# Conserved helical motifs in the Ikaros IDR mediate NuRD interaction and transcriptional repression

**DOI:** 10.1101/2024.02.29.582782

**Authors:** Tianyi Zhang, Yi-Fang Wang, Alex Montoya, Nehir Nebioglu, Husayn A. Pallikonda, Radina Georgieva, Ilinca Patrascan, James WD King, Holger B. Kramer, Pavel V. Shliaha, David S. Rueda, Matthias Merkenschlager

**Affiliations:** MRC Laboratory of Medical Sciences, Institute of Clinical Sciences, Faculty of Medicine, Imperial College London, Du Cane Road W12 0HS; Section of Virology, Department of Infectious Disease, Imperial College London, Du Cane Road, London W12 0HS

## Abstract

The transcription factor IKZF1/Ikaros is essential for B cell development, and recurrently mutated in human B-ALL. Ikaros has been ascribed both activating and repressive functions via interactions with coactivator and corepressor complexes, but the relative abundance of Ikaros-associated coregulatory complexes and their contribution to Ikaros-mediated gene regulation are not well understood. To address this issue, we performed an unbiased identification of Ikaros-interacting proteins in pre-B cells, and found that Ikaros interacts overwhelmingly with corepressors and heterochromatin-associated proteins. Time-resolved analysis of transcription and chromatin state identified transcriptional repression as the immediate response to Ikaros induction. Transcriptional repression preceded transcriptional activation by several hours, and was accompanied by a rapid loss of chromatin accessibility and reduced levels of H3K27ac particularly at enhancers. Functional characterisation of intrinsically disordered regions in the Ikaros protein identified highly conserved helical motifs that mediate Ikaros association with the NuRD corepressor complex and contribute to the silencing of target genes in pre-B cells and antiproliferative functions of Ikaros in human B-ALL.

## Introduction

In multicellular organisms, lineage-specific transcription factors (TFs) orchestrate the gene expression programs necessary for differentiation and development^1^. TFs along with a diverse range of chromatin-associated coregulators interface with regulatory elements and alter the chromatin environment to either promote or suppress transcription. Ikzf1 (Ikaros) is a hematopoietic TF essential for lymphoid commitment and pre-B cell development^2-4^. Loss of function mutations in Ikaros that lead to haploinsufficiency or dominant negative effects are frequent in pre-B cell acute lymphoblastic leukemia (B-ALL) and are associated with poor patient outcomes^5-8^.

Ikaros has been ascribed activating, repressive, pioneering, and 3D genome organising functions ^9-12^. Transcriptional control by Ikaros has been linked to both the loss and gain of chromatin accessibility and epigenetic modifications at regulatory elements across the genome^9,12-15^. What remains unresolved, especially in models looking over days or developmental timelines, are what genes or regulatory regions are direct versus secondary effects of Ikaros perturbation. Ikaros has been associated with a myriad of chromatin coactivators and corepressors ^16^. Identified interactors include the NuRD (Ref 17) and Sin3 (Ref 18) HDAC complexes as well as the corepressor CtBP1 (Ref 19). Repressive histone modifications H3K9me3 and H3K27me3 have been suggested to contribute towards Ikaros-mediated transcriptional repression ^14,20^. Conversely, interaction or cooperation between Ikaros and the BRG1 BAF chromatin remodelling complex ^21,22^, the positive elongation factor P-TEFb (Ref 23,24), and the TFs Gata-1 (Ref 23) or Gfi1 (Ref 13) suggest a role for Ikaros in gene activation. The relative abundance of coregulatory complexes associated with Ikaros, and their contribution to Ikaros-mediated gene regulation have not been fully characterised, and precisely how Ikaros binding modulates the chromatin environment to activate or repress transcription is not fully clear.

We find that the Ikaros interactome is strongly biased towards corepressors over coactivators, and that transcriptional repression is the immediate response to Ikaros induction in pre-B cells. Ikaros controls target gene expression primarily through modulating the chromatin state of regulatory elements, in particular enhancers, which are rapidly attenuated by Ikaros. Regulatory elements most sensitive to Ikaros repression have especially high Ikaros motif density and are enriched for Ikaros binding, recruitment of the NuRD subunit CHD4, and rapid H3K27ac loss. In pre-B cells, gene activation occurs later than repression, is generally not associated with direct Ikaros binding and therefore likely a secondary consequence of Ikaros. We identify and characterise a conserved helical motif region in the Ikaros IDR that mediates its association with NuRD corepressor complex, contributes to the ability of Ikaros to stably silence target genes, and attenuates the proliferation of IKZF1-mutant human B-ALL cells.

### Key Findings

- Ikaros associates predominantly with corepressive chromatin complexes
- Ikaros induction leads to rapid loss of chromatin accessibility and H3K27ac at regulatory regions, especially enhancers and superenhancers
- Conserved helical motifs in Ikaros mediate NuRD-interaction and are critical for repression
- Immediate Ikaros repressed targets in pre-B cells significantly overlap with deregulated genes in Ikaros-mutated B-ALL

## Results

### Ikaros interactome is strongly biased towards corepressors over coactivators

To identify chromatin coregulators associated with Ikaros, we performed ChIP mass spectrometry of HA-tagged Ikaros in the mouse pre-B cell line B3. We identified approximately 4000 proteins in the Ikaros chromatin-mediated interaction network (Figure 1A, Supplementary Figure 1A). Among the most enriched Ikaros interactors were DNA-binding and chromatin-associated proteins. One of the most abundant interactors was Aiolos (*Ikzf3*), a member of the Ikaros TF family that heterodimerises with Ikaros^25^ (Figure 1AB). Other highly enriched B-cell TFs included Gfi1b, Lef1, Zeb2, Ebf1, Stat5a, and Runx1 (Figure 1B). Also among the top interactors were RNA-processing factors and enzymes involved in posttranslational modification of Ikaros, such as Casein Kinase II (CK2) (Ref26), Protein Phosphatase I (PP1) (Ref27), the PIAS E3 sumo ligases^28^ (Figure1B). Ikaros affinity purification MS from the nuclear soluble fraction performed without crosslinking yielded a similar list of top interactors (Supplementary Figure 1BC), suggesting that many of these are direct protein-protein interactions.

**Figure 1.**
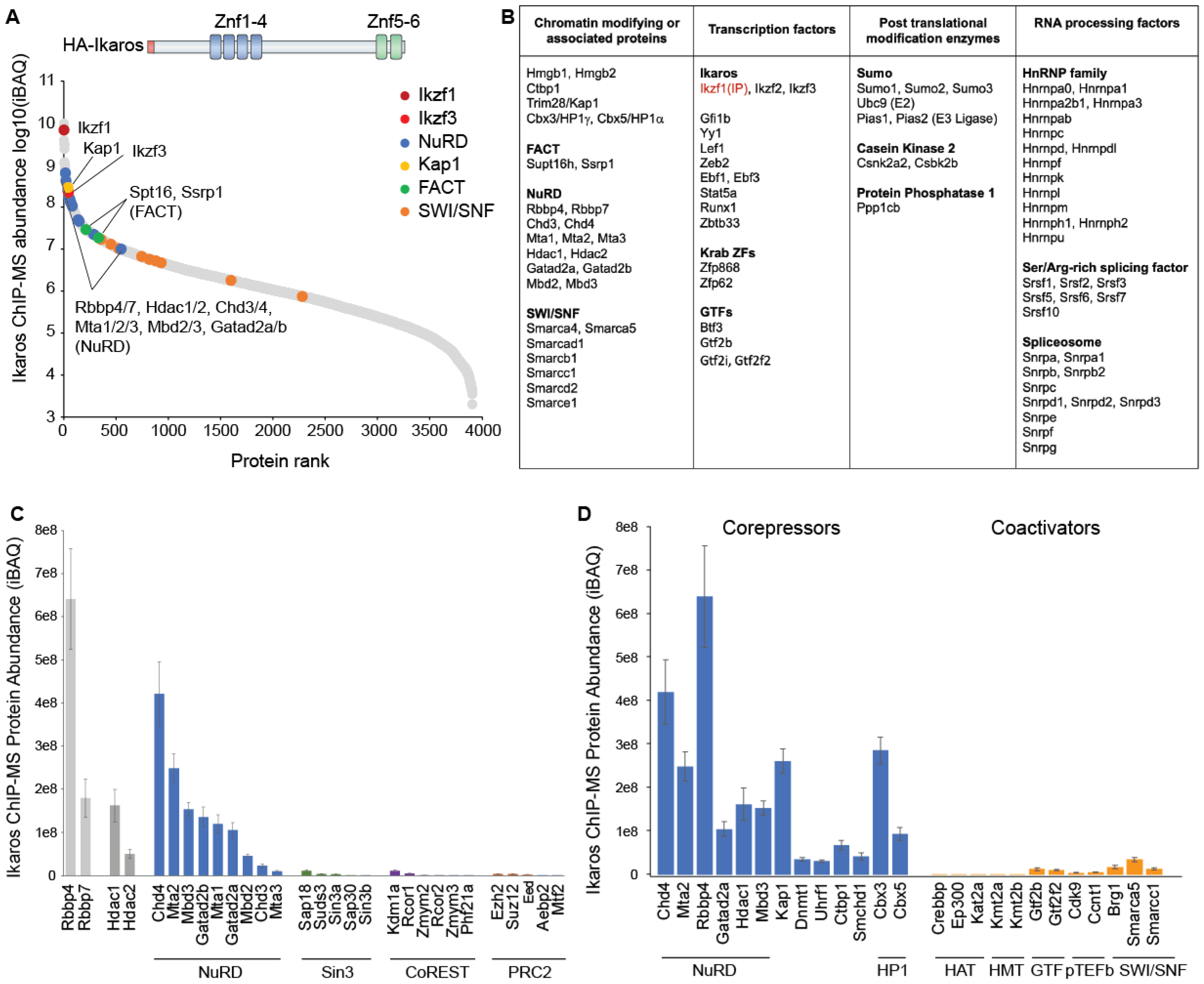
Ikaros interactome is strongly biased for corepressors over coactivators. a) Ikaros ChIP-MS interacting proteins ranked by abundance (log10 iBAQ mean of four replicates), with Ikaros (Ikzf1) in maroon, Aiolos (Ikzf3) in red, NuRD subunits in blue, Kap1 in yellow, FACT subunits in green, and SAGA SWI/SNF subunits in orange. Average of 4 replicates. b) Top 20% most abundant Ikaros ChIP-MS interacting proteins, classified into chromatin regulators, transcription factors, post-translational modification enzymes, and RNA-processing factors. c) iBAQ abundance of Hdac1/2 and Rbbp4/Rbbp7-containing complexes NuRD, Sin3, and CoREST, and the Rbbp4/7-containing complex PRC2. d) Most abundant Ikaros ChIP-MS corepressors versus coactivators.

Top interactors were associated mainly with repressive functions such as histone deacetylation, DNA methylation, and heterochromatin formation (Figure 1B). These included two known Ikaros associated corepressors CtBP1 (Ref19) and NuRD (Ref17) (Figure 1B). NuRD was the predominant Ikaros-associated HDAC complex; by comparison the Sin3 previously identified as an Ikaros interactor^18^ and CoREST were over 40-fold less abundant compared to NuRD (Figure 1C). We identified several novel Ikaros interactors associated with heterochromatin formation, DNA methylation, and H3K9 methylation including KAP1, DNMT1, UHRF1, SMCHD1, HP1α, and HP1γ (Figure 1BD). By contrast, interaction with facultative heterochromatin factors such as the PRC2 Polycomb proteins was extremely low compared to NuRD (Figure1C).

The relative association of Ikaros with corepressors was significantly more abundant than with coactivators (Figure 1D). Previously reported interactors such as the transcription elongation factor P-TEFb (Ref 23) and chromatin remodelling BRG1 SWI/SNF complex ^21^ were detected but over 10-fold less abundant than NuRD(Figure 1D). Chromatin-mediated interactions of Ikaros with coactivators p300/CBP histone acetyltransferase (HATs), H3K4 histone methyltransferases (HMTs), and the general transcription factors (GTFs) were between 20 to 2000-fold less abundant than NuRD (Figure 1D). These results reveal that in pre-B cells Ikaros overwhelmingly associates with corepressor complexes, with the relative abundance of corepressors over an order of magnitude more than interactions with coactivators.

### Transcriptional repression is the immediate response to Ikaros induction in pre-B cells

To investigate how Ikaros and associated coregulators shape the chromatin environment to regulate transcription, we employed an inducible Ikaros model using tamoxifen-induced nuclear translocation of Ikaros-ERt2 (Figure 2A). The temporal resolution afforded by this approach allowed us to identify the immediate effects of Ikaros binding on chromate state and transcription in pre-B cells.

**Figure 2.**
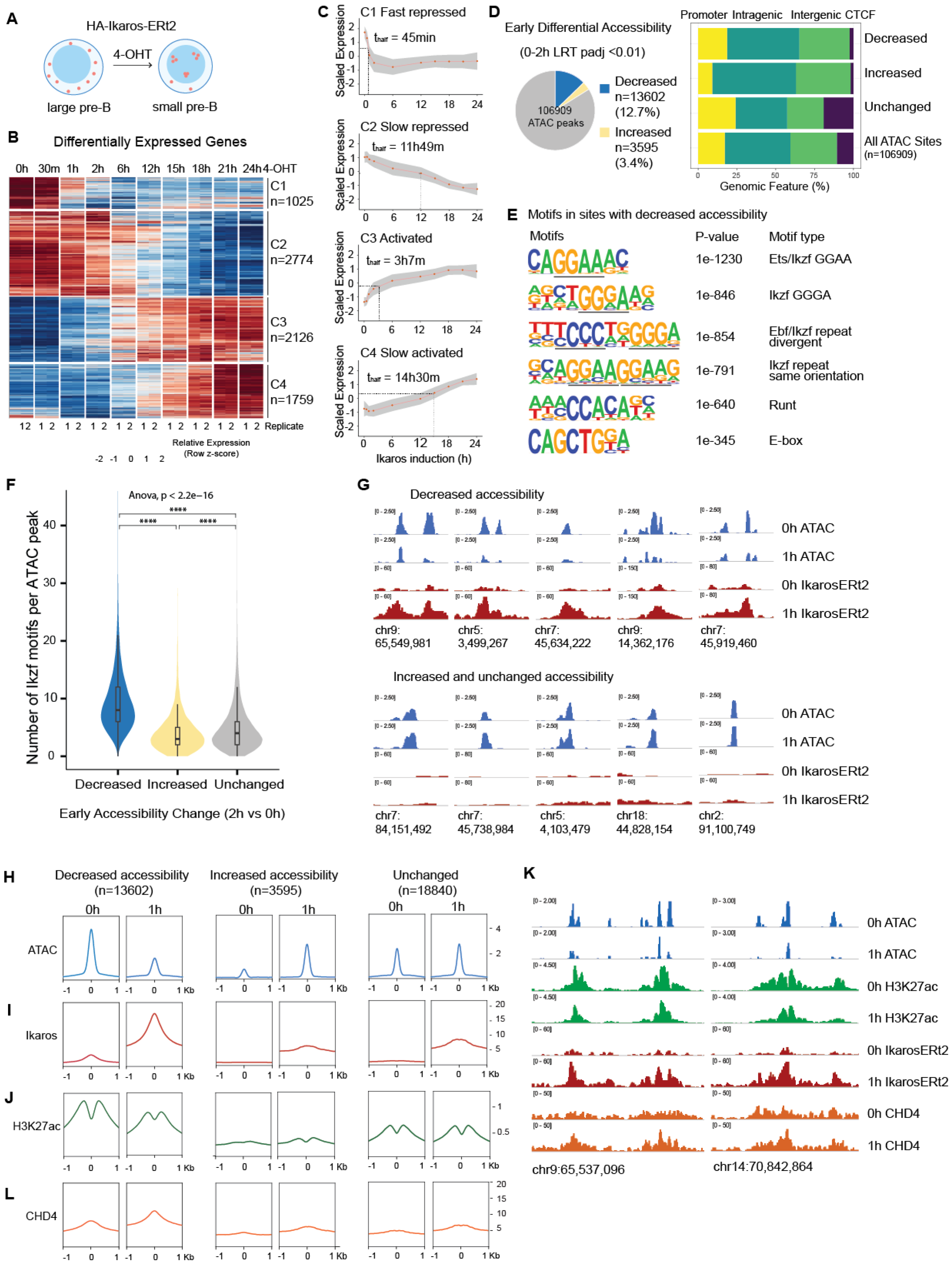
Ikaros induces rapid loss of chromatin accessibility and H3K27ac at selective sites across the genome. a) Inducible Ikaros-ERt2 pre-B model, where tamoxifen (4-OHT) induces synchronous nuclear entry of Ikaros-ERt2. b) Heatmap of differentially expressed genes (LRT padj <0.01 and log2FC>1 or <-1) over 24h nascent chromatin-associated RNA-seq timecourse following Ikaros induction performed in duplicate, and K-means clusters of genes by behaviour into C1 Fast Repressed, C2 Slow Repressed, C3 Activated and C4 Slow Activated. c) Scaled normalised expression for Kmeans clusters C1 Fast Repressed, C2 Slow Repressed, C3 Activated, and C4 Slow Activated genes, with red line and grey area indicating mean and 95% confidence intervals respectively. The time for genes in each cluster to reach half-maximal repression or activation is indicated by t-half. d) Left – Total Nucleosome Free Region (NFR) ATAC peaks called by MACS2 from ATAC-seq performed in duplicate and proportion of ATAC peaks with significant differential accessibility within 2h of Ikaros induction, decreased accessibility in blue (2h vs 0h padj<0.01, FC<0), and increased accessibility in yellow (2h vs 0h padj <0.01, FC>0). Right - Relative genomic distribution of ATAC NFR peaks at CTCF sites, Promoters, Intragenic, and Intergenic for sites with early significant decreased accessibility, early significant increased accessibility, and a subset of sites with unchanged accessibility (LRT padj>0.05, log2FC[-0.2,0.2]), and all accessible sites. e) Homer motif analysis showing top enriched motifs at ATAC NFR regions with early significant decreased accessibility compared to regions with unchanged accessibility. f) Number of Ikaros (Ikzf) motif (GGAA and GGGA) at sites with early decreased accessibility, increased accessibility, or unchanged accessibility. g) IGV browser track of ATAC and Ikaros-ERt2 at 0h and 1h following Ikaros induction at sites with early decreased accessibility (top panel) and sites with early increased or unchanged accessibility (bottom panel) h) Metaprofile plot showing chromatin accessibility at 0h and 1h following Ikaros induction at +/-1kb around sites centered at ATAC peaks with early decreased, increased or unchanged accessibility. i) Metaprofile plot showing Ikaros-ERt2 binding at 0h and 1h following Ikaros induction at +/-1kb of sites centered at ATAC peaks with early decreased, increased or unchanged accessibility. j) Metaprofile plot showing H3K27ac levels at 0h and 1h following Ikaros induction at +/-1kb of sites centered at ATAC peaks with early decreased, increased or unchanged accessibility. k) IGV browser tracks of genomic regions that show early significant decreased accessibility, with ATAC in blue, H3K27ac in green, Ikaros-ERt2 in red, and CHD4 in orange at before (0h) and 1h following Ikaros induction. l) Metaprofile plot showing CHD4 binding at 0h and 1h following Ikaros induction at regions +/-1kb of sites centered at ATAC peaks with early decreased, increased or unchanged accessibility.

Chromatin-associated-RNA-seq, which enriches for nascent transcripts, detected changes in transcription of 139 genes (126 down, 13 up) within the first 30 min of Ikaros induction. In total, 8165 genes were differentially expressed over the entire 24h timecourse (LRT padj 0.01, FC>2) (Figure 2B). K-means clustering was used to separate differentially expressed genes by behaviour (Figure 2B). Genes in clusters C1 (n=1025) and C2 (n=2774) were repressed, and genes in C3 (n=2126) and C4 (n=1759) were activated (Figure 2BC). A small group of non-monotonic genes C5 (n=481) enriched in sterol pathways linked to tamoxifen induction was removed from further analysis^29,30^ (Supplementary Figure 2AB). To determine the speed of transcriptional repression and activation, we calculated the time taken to reach half-maximal repression or activation for genes in C1-C4 (Figure 2C).

The most immediate response to Ikaros induction was rapid repression of genes in C1. Half-maximal repression of C1 genes was reached within 45 minutes of Ikaros induction, and full repression within 2h (Figure 2C). The C1 fast repressed genes were enriched in pathways related to immune processes and leukocyte differentiation, cell migration and adhesion (Supplementary Figure 2B). Notably, well characterised Ikaros regulated genes such as *Igll1* (Ref 31,32), *VpreB1* (Ref 33) and *Dntt (*Ref 34) were all found within the fast repressed class. The transcriptional response of genes in C2, C3, and C4 was slower suggesting these genes may not be primary targets of Ikaros regulation. C2 slow repressed genes were enriched for metabolic pathways and reached half their final level of repression around 12h (Figure 2C, Supplementary Figure 2B). C3 activated and C4 slow activated genes were involved in immunoglobulin production and defence response pathways reached half their final level of activation at 3h and 14.5h respectively (Figure 2C, Supplementary Figure 2B). This indicates that the immediate consequence of Ikaros induction is transcriptional repression, which precedes transcriptional upregulation by 2.5 hours or more.

### Ikaros binding drives rapid loss of chromatin accessibility and H3K27ac at regulatory promoters and enhancers

To understand how rapid transcriptional repression is accomplished, we monitored Ikaros binding and Ikaros-mediated changes to the regulatory chromatin landscape in the first 2h of induction. We used ATAC-seq and ChIP-seq for histone H3K27 acetylation to map changes in chromatin accessibility and activity of regulatory elements^35 36,37^. Ikaros induction led to rapid reduction of chromatin accessibility at a subset of accessible sites across the genome. Using ATAC-seq reads corresponding to nucleosome-free regions (NFR) which are enriched for TF binding sites^38^, we called a total of 106909 ATAC peaks (Figure 2D left panel). Within 2h of Ikaros induction, 12.7% (13602) of sites significantly decreased in accessibility while 3.4% (3595) of sites significantly increased in accessibility (LRT paj<0.01) (Figure 2D left panel). Early differentially accessible sites overlapped with promoters as well as intragenic and intergenic regions (Figure 2D right panel). Ikaros induction did not affect the accessibility of CTCF insulator sites which were depleted from early differentially accessible sites (Figure 2D right panel).

Sites with an early decrease in accessibility were highly enriched for Ikaros motifs (Figure 2E), and contained a significantly higher number and density of Ikaros motifs compared to sites with early increased and unchanged accessibility (Figure 2F, Supplementary Figure 2C). In agreement with the motif analysis, Ikaros ChIP-seq showed that Ikaros binding was strongly enriched at sites with early decreased chromatin accessibility but not at sites with early increased and unchanged accessibility (Figure 2GHI). Notably, the Ikaros peak summit was precisely centred at ATAC peaks with early decreased accessibility, which was not observed for sites with increased or unchanged accessibility where binding was more level across the region (Figure 2GHI). Sites with early increased accessibility were enriched for other TF motifs such as E-box, KLF, Forkhead, and Runt (Supplementary Figure 2D) indicating that early increased accessibility was independent of Ikaros binding.

The majority (68%) of sites with early decreased accessibility overlapped H3K27ac peaks, a signature of active promoters and enhancers (Supplementary Figure 2E). These sites showed a marked loss in H3K27ac over the flanking region in response to Ikaros binding (Figure 2JK, Supplementary Figure 2F). These data support a mechanism of repression where Ikaros drives rapid loss of focal accessibility at the site of binding, followed by a reduction in H3K27ac across the broader region.

### NuRD is recruited early to Ikaros-repressed sites

The speed and extent of decreased chromatin accessibility and H3K27ac in response to Ikaros induction suggested a process of active chromatin remodelling and deacetylation. To explore the underlying mechanisms, we focused on NuRD and KAP1, the two most abundant Ikaros-associated corepressors we identified by ChIP-MS. The NuRD complex combines chromatin remodelling and histone deacetylase activity, while KAP1 has been found to silence genes by inducing the spread of the repressive histone modification H3K9me3 (Ref39). We performed ChIP-seq of CHD4, the chromatin remodelling subunit of the NuRD complex, and found that binding of CHD4 increased after Ikaros induction and closely mirrored the binding of Ikaros itself (Figure 2IKL). As observed for Ikaros, CHD4 peaks were precisely centred and most enriched at sites with early decreased accessibility (Figure 2KL). Unlike Ikaros and CHD4 which bound strongly at accessible chromatin sites, KAP1 and H3K9me3 peaks were depleted from accessible sites (Supplementary Figure 2GH). KAP1 and H3K9me3 signal overlapped with each other within gene bodies, but unlike CHD4 KAP1 and H3K9me3 were not enriched at regions with early decreased accessibility after Ikaros induction (Supplementary Figure 2GH).

Together, these data suggest that rapid loss of chromatin accessibility and H3K27ac from active regulatory elements in response to Ikaros induction is associated with early recruitment of NuRD, but is not dependent on the acquisition of KAP1 or H3K9me3.

### Enhancers are highly sensitive to Ikaros-mediated repression

Having found that Ikaros and NuRD induce rapid repression of chromatin state at regulatory elements, we next focused on the role of promoters and enhancers in Ikaros-mediated repression. Genome-wide, enhancers were twice as likely to lose accessibility as promoters. Within 2h of Ikaros induction, 30% of accessible sites within active enhancers showed a significant decrease in chromatin accessibility compared with 15% of accessible sites within active promoters (Supplementary Figure 3A). While the levels of Ikaros and CHD4 binding were similar at active enhancers and promoters with early decreased accessibility (Figure 3AB), enhancers lost accessibility faster than promoters, and the loss of accessibility was greater at enhancers than at promoters. Differences between promoters and enhancers were even more pronounced for the loss of H3K27ac (Figure 3C).

**Figure 3.**
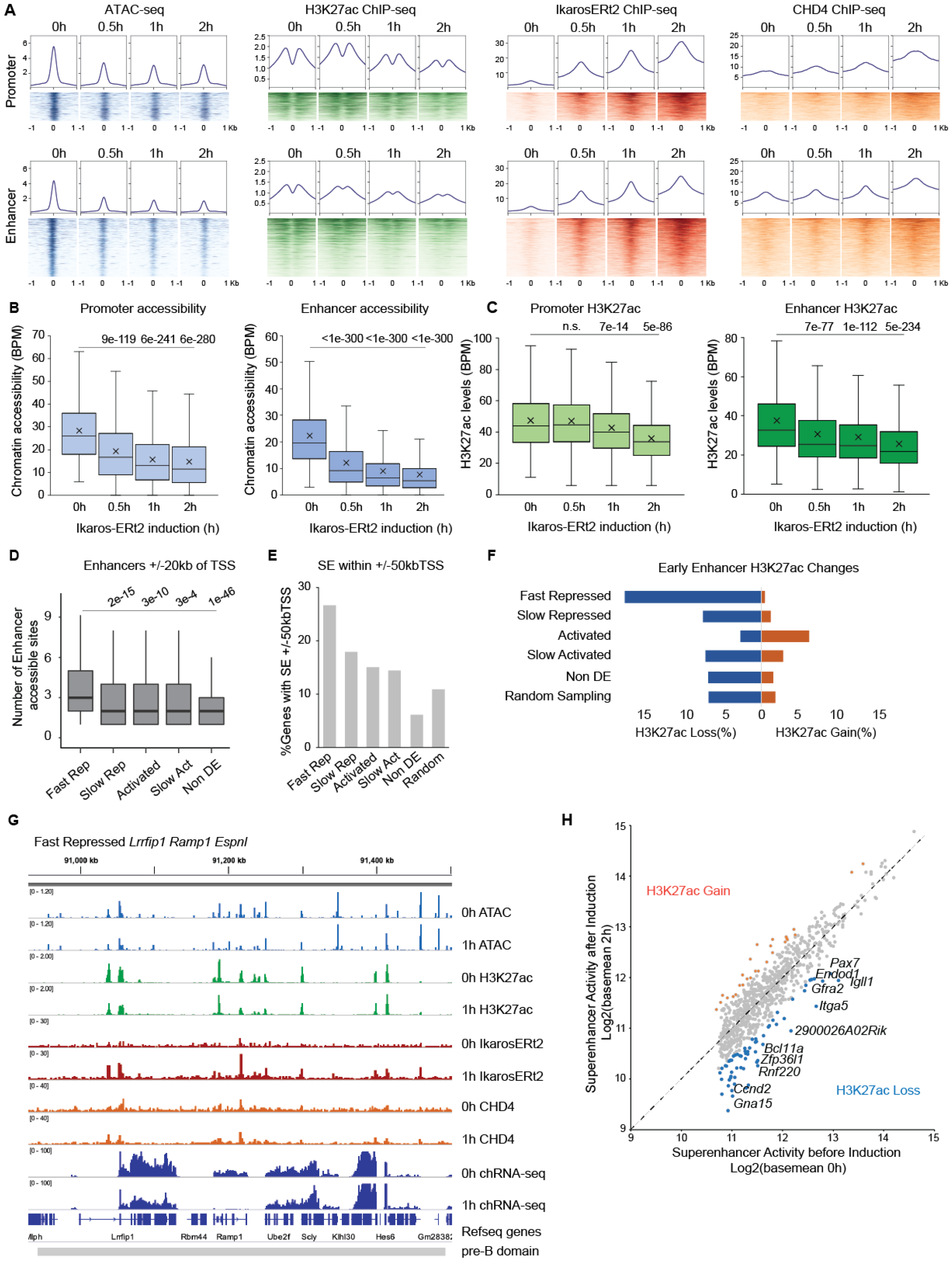
Ikaros suppresses enhancer activity to repress transcription. a) Heatmap and metaprofile plot showing chromatin accessibility, H3K27ac, Ikaros-ERt2, and CHD4 at 0h, 30m, 1h, and 2h following Ikaros induction at +/-1kb around ATAC sites with early decreased accessibility overlapping active promoters (n=2420) or enhancers (n=6273). b) Boxplot of chromatin accessibility active promoters and enhancers before (0h) and at early timepoints following Ikaros induction (30m, 1h, 2h). Pairwise student’s t-test was performed for 0h vs 30m, 1h, or 2h with Bonferroni corrected p-adjusted value stated. c) Boxplot of H3K27ac active promoters and enhancers before (0h) and at early timepoints following Ikaros induction (30m, 1h, 2h). Pairwise student’s t-test was performed for 0h vs 30m, 1h, or 2h with Bonferroni corrected p-adjusted value stated. d) Number of accessible peaks within active enhancers in the region +/-20kb TSS of Fast Repressed, Slow Repressed, Activated, Slow Activated, and non-differentially expressed genes. Pairwise Wilcoxon rank sum test was performed for Fast Repressed vs other gene class with Bonferroni corrected p-adjusted values stated. e) Proportion of genes that overlap with a SE within +/-50kb of the TSS, for Fast Repressed, Slow Repressed, Activated, Slow Activated, non-differentially expressed, and a randomly sampled set of genes. f) Proportion of enhancer H3K27ac peaks within +/-20kb of TSS Fast Repressed, Slow Repressed, Activated, Slow Activated, non-differentially expressed, and randomly sampled genes that show significant (padj<0.01, FC>1.5) loss or gain of H3K27ac within 2h of Ikaros induction. g) IGV browser of locus with three Fast Repressed genes *Lrrfip1, Ramp1, Espnl* with tracks at 0h and 1h after Ikaros induction for ATAC in blue, H3K27ac in green, Ikaros-ERt2 in red, CHD4 in orange, chRNA-seq and Refseq genes in navy, and pre-B cell domains in grey. h) Levels of H3K27ac at all SE before and 2h after Ikaros induction, superenhancers with greater than 1.5-fold increase in H3K27ac and padj <0.01 in orange, and greater than 1.5-fold decrease and padj <0.01 in H3K27ac in blue. Fast Repressed genes are labelled next to their closest SE.

H3K27ac at active enhancers decreased significantly within 30 minutes of Ikaros induction, whereas H3K27ac loss at promoters was not observable until 1h (Figure 3C). Moreover, sites with the greatest reduction in H3K27ac were found more often at enhancers than at promoters (Supplementary Figure 3b). These data show that Ikaros represses enhancer accessibility and H3K27ac more rapidly and to a greater extent than promoters.

### Preferential association of fast repressed genes with enhancers

Consistent with a model where enhancers are primary targets of Ikaros-mediated repression, we found that fast repressed genes were associated with a higher local density of enhancers within 20kb (Figure 3D) and a greater number of Ikaros-bound sites at enhancers within 20kb (Supplementary Figure 3C). A greater proportion of fast repressed genes (26%) were associated with superenhancers (SEs) compared to slow responding and non-differentially expressed genes (Figure 3E). Additionally, to assign enhancers to promoters using a more data-driven approach, we used Fantom5 CAGE-seq to correlate enhancer and gene transcription in combination with Hi-C contact frequency information (Methods). This analysis further confirmed that Fast Repressed genes had more numerous enhancer associations than other gene classes (Supplementary Figure 3D).

Not only were fast repressed genes associated with more enhancers overall, these enhancers were also more sensitive to Ikaros-mediated repression. Enhancers associated with fast repressed genes showed a greater loss in both chromatin accessibility and H3K27ac within the first 2h of Ikaros induction compared to enhancers associated with other groups (Figure 3F, Supplementary Figure 3E).

Enhancers proximal to fast repressed genes were approximately twice as likely to lose accessibility and three times more likely to lose H3K27ac compared with enhancers near randomly sampled genes (Supplementary Figure 3FG). By comparison, enhancers associated with slow repressed, activated, slow activated and non-differentially expressed genes were less likely to lose accessibility and H3K27ac within 2h of Ikaros induction (Figure 3F, Supplementary Figure 3FG). We observed many examples of fast repressed genes where Ikaros binding at nearby enhancers induced rapid loss of accessibility and H3K27ac (Supplementary Figure 3H), and multiple fast repressed genes were in the vicinity of repressed enhancers (Figure 3G).

Finally, we saw a strong effect of Ikaros-mediated repression on SE activity. More than half of SEs (56%) that showed a significant early decrease in H3K27ac were associated with a fast repressed gene. Figure 3H shows examples of several fast repressed genes including *Itga5, Endod1, Ccnd2, Igll1, Bcl11a*, and *Gfra2* associated with SEs showing more than 1.5-fold decrease in H3K27ac by 2h of Ikaros induction. These results together with the observation that enhancers show greater sensitivity to Ikaros induction, suggest that enhancer and SE disruption may be a primary driver of Ikaros-mediated transcriptional repression.

### Identification of conserved helical motifs in Ikaros IDR

To determine domains within the Ikaros protein that confer repressive function we focused on intrinsically disordered regions (IDRs). As is the case for most TFs, large parts of Ikaros, the N-terminal region and the internal region between ZnF4 and ZnF5, are unstructured (Figure 4A, Supplementary Figure 4A). The PEST regions (proline, glutamic acid, serine, threonine-rich sequences) are known to regulate Ikaros protein stability^27^, but otherwise the functional significance of Ikaros IDRs remains poorly defined. Using the 9aaTAD prediction tool which identifies conserved alpha-helical motifs present in many diverse eukaryotic TFs^40^, we identified three sequences in Ikaros strongly matching the 9aaTAD pattern (Figure 4BC, Supplementary Figure 4B)^40,41^. All three were located in the internal Ikaros IDR within two alpha-helices in the Ikaros Alphafold structure (Figure 4D, Supplementary Figure 4A). The three motifs, SLVLDRLAS in Helix 1 and DMMTSHVMD and QAINNAINY in Helix 2, as well as several residues following both helices are highly conserved between Ikaros family TFs (Figure 4C).

**Figure 4.**
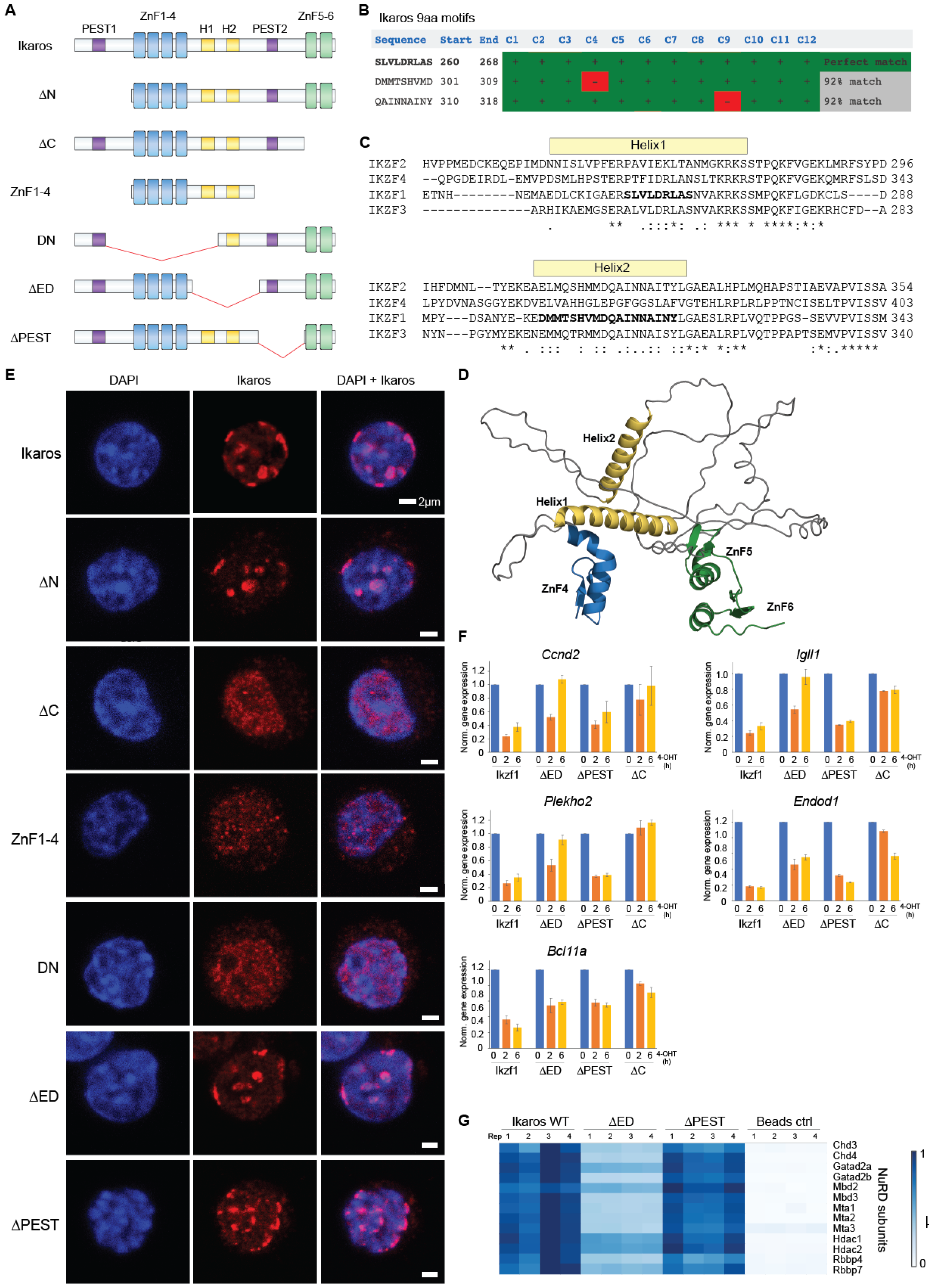
conserved helical motifs in the Ikaros IDR is required for gene repression and NuRD interaction. a) Schematic of Ikaros and truncation mutants, ΔN (N-terminus IDR deletion), ΔC (C-terminus ZnF 5-6 deletion), Znf1-4 (DNA-binding ZnF1-4), DN (leukemia associated dominant negative isoform ik6), ΔED (effector domain deletion), ΔPEST (proline, glutamic acid, serine, threonine-rich domain deletion) b) Top three 9aa motifs in the mouse Ikaros protein obtained from the 9aaTAD Prediction Tool. c) Protein sequence alignment of the IDR region of mouse Ikaros IKZF1 with close paralogs IKZF2, IKZF3 and IKZF4, with the three predicted 9aa motifs in bold, position of alpha helices shown in yellow and Clustal Omega amino acid conservation scores shown below the alignment. d) Alphafold structure of Ikaros, showing the internal disordered region between ZnF4 (blue) and ZnF5 (green), with the conserved helices containing the 9aa motifs in yellow. e) Immunofluorescence of B3 cells expressing HA-tagged full length Ikaros and truncation mutants ΔN, ΔC, ZnF1-4, DN, ΔED, and ΔPEST. f) qPCR of normalized gene expression at several Fast Repressed genes before and 2h and 6h after induction with wildtype Ikaros or mutants ΔED, ΔPEST and ΔC. g) Heatmap of NuRD complex subunit abundance in affinity purification-MS of Ikaros, ΔED, ΔPEST, and untransduced cells.

We generated a truncation mutant of the conserved helical motifs (effector domain deletion ΔED) and first assessed whether this region was necessary for DNA-binding. Ikaros forms foci at pericentric major satellite repeats which contain the Ikaros binding motif GGAA. These foci serve as indicators of Ikaros DNA binding activity, which is abrogated by removal of the DNA-binding (Dominant-negative DN) or dimerisation domain (ΔC) (Figure 4E) as previously shown ^42,43^. By contrast, removal of the conserved helices in ΔED, or in fact any of the IDR regions ΔN, ΔED, or ΔPEST, had no impact on DNA-binding in B3 cells (Figure 4E).

### Conserved helical motifs mediate transcriptional repression and NuRD interaction

The region overlapping Helix2 was previously identified as a transactivation domain in yeast-one-hybrid assays ^44^, but did not exhibit activation function in cellular reporter assays ^45^. We thus asked whether the helical motifs were involved in transcriptional repression. We assayed the ability of the ΔED mutant to repress several Ikaros fast repressed genes we identified. Induction of wild-type Ikaros resulted in silencing of fast repressed genes, which was complete by 2h and sustained at 6h (Figure 4f, Supplementary Figure 4C). The ΔC mutant which cannot repress due to defective DNA-binding served as a negative control (Figure 4F, Supplementary Figure 4C). Ikaros ΔED (which lacked the helical motifs) showed weak repressive ability at 2h, but this repression was not sustained at 6h (Figure 4F). Deletion of the conserved residues containing Helix1 or Helix2 in isolation (mutants ΔED1 and ΔED2) showed partial silencing defects, suggesting that the entire conserved helical region is important for the repressive ability of Ikaros (Supplementary F4CDE). We also assayed another IDR mutant ΔPEST which contains the conserved helices but lacked the adjacent PEST-sequence. In contrast to the ΔED mutant, the ΔPEST mutant showed near wild-type repression of Ikaros target genes which supports a role for the conserved helical motifs in Ikaros-mediated repression (Figure 4F).

As Ikaros IDR mutants had normal DNA-binding function, we asked whether the reduced repressive function of ΔED was due to loss of a protein-protein interaction mediated through the conserved helical motifs. To this end, we performed affinity purification MS of wild-type Ikaros, the silencing-defective ΔED, the silencing-competent ΔPEST. Strikingly, while most interactors were preserved between Ikaros ΔED and wild-type, ΔED showed substantially reduced interactions with all NuRD complex subunits (Figure 4G, Supplementary Figure 4F). By contrast, ΔPEST retained wild-type levels of NuRD association (Figure 4G, Supplementary 4F). These data demonstrate that the conserved helical motifs within the internal IDR of Ikaros mediate the repressive function of Ikaros through interaction with the NuRD complex.

### Conserved helical motifs are required for the anti-proliferative function of Ikaros

We next investigated the contribution of the conserved helical motifs to Ikaros function in human B-ALL cells, as evolutionary conservation of this region across all vertebrates suggested a functional importance that is also likely to be conserved in humans. Ikaros acts as a tumour suppressor in acute pre-B and T cell leukemia, and exhibits anti-proliferative effects through repression of *Myc* and cell cycle genes^46^. In B-ALL, recurring deletions of the DNA-binding domain in Ikaros generate the dominant negative Ik6 (DN Ik6) mutant associated with the aggressive BCR-ABL1 subtype^5,8^. As found in previous studies^47,48^ we found that expression of Ikaros reduced the growth of leukemic cell lines. In B-ALL lines SupB15 and BV173 which express high levels of DN Ikaros mutant, we found that forced expression of exogenous wild-type Ikaros reduced the growth rate of these cells (Figure 5AB). Importantly, this anti-proliferative effect was reduced or absent in the ΔED Ikaros mutant (Figure 5CD). This indicates that the conserved helical motif region is required for anti-proliferative functions of Ikaros in human B-ALL cells. In agreement with these data, missense mutations in this region of Ikaros, specifically in residues in Helix1, Helix2, and the conserved stretch after Helix2, have been identified in patients with adult T-ALL^49^ and in germline mutations of pediatric B-ALL^50^ (Figure 5E). This suggests that disruption of residues in this region of Ikaros which confers NuRD binding and repressive function could contribute to leukemogenesis.

**Figure 5.**
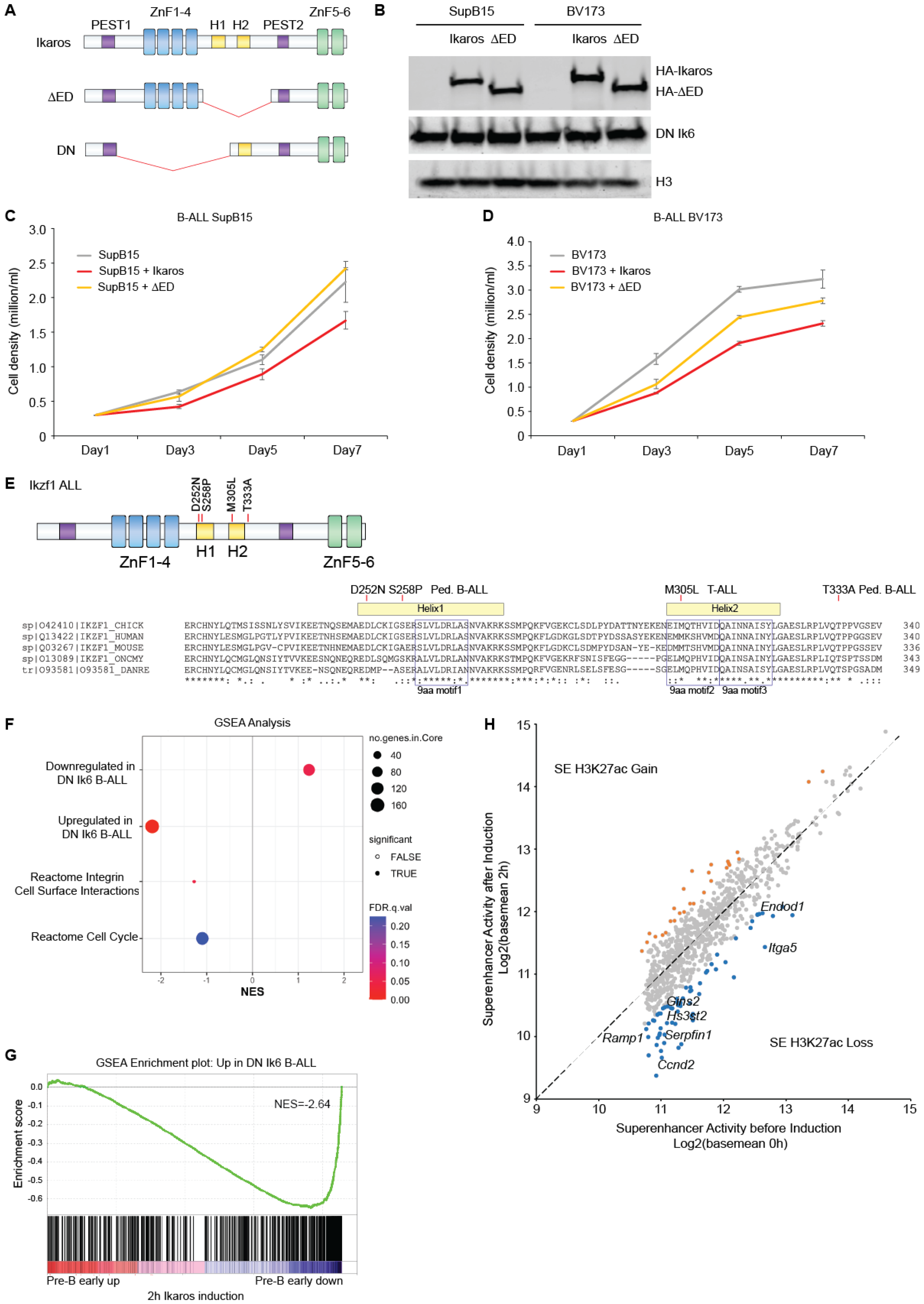
Conserved helical motifs are required for the anti-proliferative function of Ikaros in B-ALL cells. a) Schematic of Ikaros, ΔED, and leukemic Ikaros dominant negative DN Ik6 mutant. b) Western blot of human B-ALL lines SupB15 and BV173 with mutated Ikaros dominant negative Ik6 transduced with Ikaros or ΔED. c) Growth curve of human B-ALL line SupB15, and SupB15 transduced with Ikaros or ΔED. d) Growth curve of human B-ALL line BV173, and BV173 transduced with Ikaros or ΔED. e) Missense mutations in the conserved helical motif region of Ikaros detected in patients with T-ALL or pediatric B-ALL, and Clustal Omega alignment of Ikaros sequence conservation between vertebrate species chicken, human, mouse, rainbow trout (*Oncorhynchus mykiss*), and zebrafish (*Danio rerio*). f) GSEA analysis of early (2h) Ikaros-induced transcriptional changes in pre-B cells with gene sets for deregulated genes in DN Ik6-expressing B-ALL, integrin and cell cycle pathways, with NES, significance, and FDR values indicated. g) GSEA enrichment plot of early (2h) Ikaros-induced transcriptional changes in pre-B cells compared with gene set of aberrantly upregulated genes in DN Ik6-expressing B-ALL. h) Superenhancer associated Fast Repressed genes aberrantly upregulated in Ikaros DN mutant B-ALL, and the degree of H3K27ac change 2h after Ikaros induction, where superenhancers with greater than 1.5-fold increase in H3K27ac and padj <0.01 in orange, and greater than 1.5-fold decrease and padj <0.01 in H3K27ac in blue.

### Deregulation of Fast Repressed genes in Ikaros-mutated leukemia

Finally, we examined the relationship between Ikaros-repressed pre-B cell genes and deregulated pathways in Ikaros-mutated leukemia. A previous model of BCR-ABL1 B-ALL found that disruption of Ikaros function through Ikaros DN mutant expression results in marked changes in the formation of sphere-like aggregates and increased adherence, with increased expression of adhesion, self-renewal, and stem-cell genes^51^. We also saw cell cycle and integrin gene sets enriched at early 2h Ikaros-repressed genes in pre-B cells (Figure 5F). Strikingly, we found substantial overlap between genes deregulated in Ikaros DN mutant B-ALL and early Ikaros repressed genes reported here (Figure 5FG). 20% of genes (101/512) found aberrantly upregulated in Ikaros DN B-ALL corresponded to genes in the Ikaros fast repressed group. We saw that several fast repressed genes upregulated in Ikaros DN mutant B-ALL including *Itga5, Endod1, Ramp1, Serpinf1, Gins2, Hs3st2 and Hs3st3b1* were associated with early Ikaros-repressed pre-B superenhancers (Figure 5H). Taken together, these results show that a number of genes we identify as direct targets of early repression by Ikaros and NuRD in pre-B cells are critically dependent on Ikaros for their repression, and become aberrantly overexpressed in Ikaros-mutated leukemia.

## Discussion

Studies from many labs have established Ikaros as a key regulator of lymphocyte development^2-4^, and as an important tumour suppressor gene in B-ALL^5-8^. Based on time-resolved analyses, we find that the immediate responses to Ikaros binding in pre-B cells are loss of chromatin accessibility and H3K27ac at regulatory enhancers, and transcriptional repression. Accordingly, proteomic quantification of Ikaros-associated proteins in pre-B cells showed that corepressive chromatin complexes by far outweigh interactions with coactivators. Dissection of intrinsically disordered regions identified highly conserved helical motifs that mediate the interaction between Ikaros and the NuRD corepressor complex, contribute to target gene silencing, and attenuate the proliferation of Ikaros mutant human B-ALL cells.

Focusing on the early direct impact of Ikaros on chromatin regulation, we observed that enhancers exhibited faster chromatin state changes in response to Ikaros than promoters. Fast repressed genes were highly enriched in pathways related to immune differentiation, and were associated with a greater number of enhancers.

Our study agrees with previous reports that hematopoietic genes associated with a greater number of enhancers and superenhancers exhibited faster expression changes during differentiation^52^, and extends these data by defining a direct role for Ikaros and NuRD in enhancer regulation through repression of the chromatin state.

We identified a conserved helical motif region in the internal IDR of Ikaros that is critical for its repressive function and for NuRD binding. The residues KRKSSMPQ in at the end Ikaros Helix1 and adjacent loop match a 12 amino acid motif present in other TFs known to interact with NuRD including Fog1, Bcl11a, and Sall1-4^53-56^ (Supplementary Figure 5AB). This region was identified as being present in Ikaros family proteins across all deuterostome phyla^57^, but had not been functionally characterised. This (R/K)RKXXXPQ motif in Ikaros and the NuRD-interacting TFs Fog1, Bcl11a, and Sall4 binds RBBP5 and resembles the interaction between the histone H3 tail with RBBP4^58^. Prediction modelling by AlphaPullDown confirmed this putative interaction between this (R/K)RKXXXPQ motif in Ikaros with RBBP4 with a quality ipTM score of 0.73 (Supplementary Figure 5C). Similar to what is observed in the crystal structures of Fog1 and Bcl11a with RBBP4, the positively charged residues “KRK” at the end of Ikaros Helix1 insert into the negatively charged central pocket of RBBP4^53-55^ (Supplementary Figure 5DE). The flexible loop that follows Helix1 interacts with the groove between two blades of the beta-propeller of RBBP4 in a fashion almost identical to that observed for H3 and the NuRD-interacting TFs^53-55,58^. An interesting difference is the (R/K)RKXXXPQ motif is N-terminal in Fog1, Bcll11a, and Sall4, but lies in the internal IDR of all Ikaros family members.

Sumoylation sites are a common feature of TF repression domains^59^, and sumoylation of Ikaros can disrupt interactions with NuRD^28^. Lys240 in Ikaros which is close to the conserved Helix1 motif identified here is one of three sumoylation sites in Ikaros^28,60^. This raises the strong possibility that the sumoylation to the conserved helical motifs identified here may be targeted by post-translational mechanisms which affect the ability of Ikaros to interact with NuRD and thus its transcriptional repression activity. Of note, our functional characterisation of Ikaros mutants suggests that the conserved region around Helix2 is similarly required for full repressive function of Ikaros, and our proteomics data indicates that Ikaros interacts significantly more strongly with NuRD than with other RBBP4-containing complexes. It is therefore likely that the putative interaction between Helix1 residues and RBBP4 represents only one of several interaction interfaces between Ikaros and NuRD.

With respect to the role of Ikaros in human disease, we find substantial overlap between early repressed genes identified here and aberrantly upregulated genes in Ikaros-mutated B-ALL. Moreover, several Ikaros-repressed super-enhancer-associated genes *ITGA5, CCND2, GFRA2, BCL11A* have been implicated in multiple cancers^61-65^. This indicates that a large number of genes aberrantly expressed in Ikaros-mutated leukemia are highly sensitive to Ikaros dosage, and dependent on Ikaros and NuRD for repression. The molecular insights uncovered for Ikaros-mediated transcriptional repression in pre-B cells are therefore likely to inform potential mechanisms of deregulation in leukemia.

## Acknowledgements

We thank LMS staff scientists Georgia Roumelioti for assistance with proteomic sample processing, Ivan Andrew and Laurence Game with sequencing, James Elliott and Bhavik Patel for cell sorting. We thank Markus Muschen and Niklas Feldhahn for the gift of B-ALL lines. We acknowledge Taha Shahid and Antoine Hocher for help with AlphaPullDown. We are grateful to members of the Lymphocyte Development, Single Molecule Imaging and Epigenetic Memory labs for helpful discussions, especially Paul Girvan for critical reading of the manuscript. This work was supported by the Medical Research Council, Wellcome Trust Grant (215933/Z/19/Z to TZ), and the Leverhulme Research Grant (RPG-2016-214 to DSR and MM).

## Author Contributions

TZ and MM conceived the experiments. TZ performed the experiments and generated the data. YW and AM made significant contributions to the bioinformatics and proteomics data analysis. NN, HAP, RG, JWDK, IP, HBK, PVS, DSR contributed to data analysis, data interpretation, or experimental design. TZ wrote the manuscript with significant input from MM, with revisions from DSR and contributions to the methods section from YW, AM and RG. DSR and MM supervised the study.

## Data availability

NGS data generated for this study have been deposited at GEO under accession number GSE256299.

## References

1 Ong, C. T. & Corces, V. G. Enhancer function: new insights into the regulation of tissue-specific gene expression. Nat Rev Genet 12, 283–293, doi:10.1038/nrg2957 (2011).

2 Georgopoulos, K. et al. The Ikaros gene is required for the development of all lymphoid lineages. Cell 79, 143–156, doi:10.1016/0092-8674(94)90407-3 (1994).

3 Lo, K., Landau, N. R. & Smale, S. T. LyF-1, a transcriptional regulator that interacts with a novel class of promoters for lymphocyte-specific genes. Mol Cell Biol 11, 5229–5243, doi:10.1128/mcb.11.10.5229-5243.1991 (1991).

4 Georgopoulos, K., Moore, D. D. & Derfler, B. Ikaros, an early lymphoid-specific transcription factor and a putative mediator for T cell commitment. Science 258, 808–812, doi:10.1126/science.1439790 (1992).

5 Mullighan, C. G. et al. BCR-ABL1 lymphoblastic leukaemia is characterized by the deletion of Ikaros. Nature 453, 110–114, doi:10.1038/nature06866 (2008).

6 Nakase, K. et al. Dominant negative isoform of the Ikaros gene in patients with adult B-cell acute lymphoblastic leukemia. Cancer Res 60, 4062–4065 (2000).

7 Martinelli, G. et al. IKZF1 (Ikaros) deletions in BCR-ABL1-positive acute lymphoblastic leukemia are associated with short disease-free survival and high rate of cumulative incidence of relapse: a GIMEMA AL WP report. J Clin Oncol 27, 5202–5207, doi:10.1200/JCO.2008.21.6408 (2009).

8 Mullighan, C. G. et al. Genome-wide analysis of genetic alterations in acute lymphoblastic leukaemia. Nature 446, 758–764, doi:10.1038/nature05690 (2007).

9 Ding, Y. et al. Ikaros tumor suppressor function includes induction of active enhancers and super-enhancers along with pioneering activity. Leukemia 33, 2720–2731, doi:10.1038/s41375-019-0474-0 (2019).

10 Lemarie, M., Botardi, S., Mavoungou, L., Pak, H. & Milot, E. IKAROS is required for the measured response of NOTCH target genes upon external NOTCH signaling. PLoS Genet 17, e1009478, doi:10.1371/journal.pgen.1009478 (2021).

11 Liang, Z. et al. A high-resolution map of transcriptional repression. Elife 6, doi:10.7554/eLife.22767 (2017).

12 Hu, Y. et al. Lineage-specific 3D genome organization is assembled at multiple scales by IKAROS. Cell 186, 5269–5289 e5222, doi:10.1016/j.cell.2023.10.023 (2023).

13 Sun, W. et al. GFI1 Cooperates with IKZF1/IKAROS to Activate Gene Expression in T-cell Acute Lymphoblastic Leukemia. Mol Cancer Res 20, 501–514, doi:10.1158/1541-7786.MCR-21-0352 (2022).

14 Song, C. et al. Epigenetic regulation of gene expression by Ikaros, HDAC1 and Casein Kinase II in leukemia. Leukemia 30, 1436–1440, doi:10.1038/leu.2015.331 (2016).

15 Schjerven, H. et al. Genetic analysis of Ikaros target genes and tumor suppressor function in BCR-ABL1(+) pre-B ALL. J Exp Med 214, 793–814, doi:10.1084/jem.20160049 (2017).

16 Affar, M. et al. IKAROS: from chromatin organization to transcriptional elongation control. Cell Death Differ, doi:10.1038/s41418-023-01212-2 (2023).

17 Sridharan, R. & Smale, S. T. Predominant interacotin of both Ikaros and Helios with the NuRD complex in immature thymocytes. J Biol Chem 282, 30227–30238, doi:10.1074/jbc.M702541200 (2007).

18 Koipally, J., Renold, A., Kim, J. & Georgopoulos, K. Repression by Ikaros and Aiolos is mediated through histone deacetylase complexes. EMBO J 18, 3090–3100, doi:10.1093/emboj/18.11.3090 (1999).

19 Koipally, J. & Georgopoulos, K. Ikaros interactions with CtBP reveal a repression mechanism that is independent of histone deacetylase activity. J Biol Chem 275, 19594–19602, doi:10.1074/jbc.M000254200 (2000).

20 Oravecz, A. et al. Ikaros mediates gene silencing in T cells through Polycomb repressive complex 2. Nat Commun 6, 8823, doi:10.1038/ncomms9823 (2015).

21 O’Neill, D. W. et al. An ikaros-containing chromatin-remodeling complex in adult-type erythroid cells. Mol Cell Biol 20, 7572–7582, doi:10.1128/MCB.20.20.7572-7582.2000 (2000).

22 Bossen, C. et al. The chromatin remodeler Brg1 activates enhancer repertoires to establish B cell identity and modulate cell growth. Nat Immunol 16, 775–784, doi:10.1038/ni.3170 (2015).

23 Botardi, S. et al. Ikaros interacts with P-TEFb and cooperates with GATA-1 to enhance transcription elongation. Nucleic Acids Res 39, 3505–3519, doi:10.1093/nar/gkq1271 (2011).

24 Botardi, S. et al. The IKAROS interaction with a complex including chromatin remodeling and transcription elongation activities is required for hematopoiesis. PLoS Genet 10, e1004827, doi:10.1371/journal.pgen.1004827 (2014).

25 Kelley, C. M. et al. Helios, a novel dimerization partner of Ikaros expressed in the earliest hematopoietic progenitors. Curr Biol 8, 508–515, doi:10.1016/s0960-9822(98)70202-7 (1998).

26 Gomez-del Arco, P., Maki, K. & Georgopoulos, K. Phosphorylation controls Ikaros’s ability to negatively regulate the G(1)-S transition. Mol Cell Biol 24, 2797–2807, doi:10.1128/MCB.24.7.2797-2807.2004 (2004).

27 Popescu, M. et al. Ikaros stability and pericentromeric localization are regulated by protein phosphatase 1. J Biol Chem 284, 13869–13880, doi:10.1074/jbc.M900209200 (2009).

28 Gomez-del Arco, P., Koipally, J. & Georgopoulos, K. Ikaros SUMOylation: switching out of repression. Mol Cell Biol 25, 2688–2697, doi:10.1128/MCB.25.7.2688-2697.2005 (2005).

29 Holleran, A. L., Lindenthal, B., Aldaghlas, T. A. & Kelleher, J. K. Effect of tamoxifen on cholesterol synthesis in HepG2 cells and cultured rat hepatocytes. Metabolism 47, 1504–1513, doi:10.1016/s0026-0495(98)90078-6 (1998).

30 Hultsch, S. et al. Association of tamoxifen resistance and lipid reprogramming in breast cancer. BMC Cancer 18, 850, doi:10.1186/s12885-018-4757-z (2018).

31 Sabbattini, P. et al. Binding of Ikaros to the lambda5 promoter silences transcription through a mechanism that does not require heterochromatin formation. EMBO J 20, 2812–2822, doi:10.1093/emboj/20.11.2812 (2001).

32 Thompson, E. C. et al. Ikaros DNA-binding proteins as integral components of B cell developmental-stage-specific regulatory circuits. Immunity 26, 335–344, doi:10.1016/j.immuni.2007.02.010 (2007).

33 Ferreiros-Vidal, I. et al. Genome-wide identification of Ikaros targets elucidates its contribution to mouse B-cell lineage specification and pre-B-cell differentiation. Blood 121, 1769–1782, doi:10.1182/blood-2012-08-450114 (2013).

34 Trinh, L. A. et al. Down-regulation of TDT transcription in CD4(+)CD8(+) thymocytes by Ikaros proteins in direct competition with an Ets activator. Genes Dev 15, 1817–1832, doi:10.1101/gad.905601 (2001).

35 Witkopp, P. J. & Kalay, G. Cis-regulatory elements: molecular mechanisms and evolutionary processes underlying divergence. Nat Rev Genet 13, 59–69, doi:10.1038/nrg3095 (2011).

36 Creyghton, M. P. et al. Histone H3K27ac separates active from poised enhancers and predicts developmental state. Proc Natl Acad Sci U S A 107, 21931–21936, doi:10.1073/pnas.1016071107 (2010).

37 Rada-Iglesias, A. et al. A unique chromatin signature uncovers early developmental enhancers in humans. Nature 470, 279–283, doi:10.1038/nature09692 (2011).

38 Yan, F., Powell, D. R., Curtis, D. J. & Wong, N. C. From reads to insight: a hitchhiker’s guide to ATAC-seq data analysis. Genome Biol 21, 22, doi:10.1186/s13059-020-1929-3 (2020).

39 Groner, A. C. et al. KRAB-zinc finger proteins and KAP1 can mediate long-range transcriptional repression through heterochromatin spreading. PLoS Genet 6, e1000869, doi:10.1371/journal.pgen.1000869 (2010).

40 Piskacek, S. et al. Nine-amino-acid transactivation domain: establishment and prediction utilities. Genomics 89, 756–768, doi:10.1016/j.ygeno.2007.02.003 (2007).

41 Piskacek, M., Havelka, M., Rezacova, M. & Knight, A. The 9aaTAD Transactivation Domains: From Gal4 to p53. PLoS One 11, e0162842, doi:10.1371/journal.pone.0162842 (2016).

42 Brown, K. E. et al. Association of transcriptionally silent genes with Ikaros complexes at centromeric heterochromatin. Cell 91, 845–854, doi:10.1016/s0092-8674(00)80472-9 (1997).

43 Cobb, B. S. et al. Targeting of Ikaros to pericentromeric heterochromatin by direct DNA binding. Genes Dev 14, 2146–2160, doi:10.1101/gad.816400 (2000).

44 Sun, L., Liu, A. & Georgopoulos, K. Zinc finger-mediated protein interactions modulate Ikaros activity, a molecular control of lymphocyte development. EMBO J 15, 5358–5369 (1996).

45 Koipally, J., Heller, E. J., Seavit, J. R. & Georgopoulos, K. Unconventional potentiation of gene expression by Ikaros. J Biol Chem 277, 13007–13015, doi:10.1074/jbc.M111371200 (2002).

46 Ma, S. et al. Ikaros and Aiolos inhibit pre-B-cell proliferation by directly suppressing c-Myc expression. Mol Cell Biol 30, 4149–4158, doi:10.1128/MCB.00224-10 (2010).

47 Dumortier, A. et al. Notch activation is an early and critical event during T-Cell leukemogenesis in Ikaros-deficient mice. Mol Cell Biol 26, 209–220, doi:10.1128/MCB.26.1.209-220.2006 (2006).

48 Kathrein, K. L., Lorenz, R., Innes, A. M., Griffiths, E. & Winandy, S. Ikaros induces quiescence and T-cell differentiation in a leukemia cell line. Mol Cell Biol 25, 1645–1654, doi:10.1128/MCB.25.5.1645-1654.2005 (2005).

49 Richter-Pechanska, P. et al. Identification of a genetically defined ultra-high-risk group in relapsed pediatric T-lymphoblastic leukemia. Blood Cancer J 7, e523, doi:10.1038/bcj.2017.3 (2017).

50 Churchman, M. L. et al. Germline Genetic IKZF1 Variation and Predisposition to Childhood Acute Lymphoblastic Leukemia. Cancer Cell 33, 937–948 e938, doi:10.1016/j.ccell.2018.03.021 (2018).

51 Churchman, M. L. et al. Efficacy of Retinoids in IKZF1-Mutated BCR-ABL1 Acute Lymphoblastic Leukemia. Cancer Cell 28, 343–356, doi:10.1016/j.ccell.2015.07.016 (2015).

52 Huang, J. et al. Dynamic Control of Enhancer Repertoires Drives Lineage and Stage-Specific Transcription during Hematopoiesis. Dev Cell 36, 9–23, doi:10.1016/j.devcel.2015.12.014 (2016).

53 Lejon, S. et al. Insights into association of the NuRD complex with FOG-1 from the crystal structure of an RbAp48.FOG-1 complex. J Biol Chem 286, 1196–1203, doi:10.1074/jbc.M110.195842 (2011).

54 Moody, R. R. et al. Probing the interaction between the histone methyltransferase/deacetylase subunit RBBP4/7 and the transcription factor BCL11A in epigenetic complexes. J Biol Chem 293, 2125–2136, doi:10.1074/jbc.M117.811463 (2018).

55 Liu, B. H. et al. Targeting cancer addiction for SALL4 by shifting its transcriptome with a pharmacologic peptide. Proc Natl Acad Sci U S A 115, E7119–E7128, doi:10.1073/pnas.1801253115 (2018).

56 Lauberth, S. M. & Rauchman, M. A conserved 12-amino acid motif in Sall1 recruits the nucleosome remodeling and deacetylase corepressor complex. J Biol Chem 281, 23922–23931, doi:10.1074/jbc.M513461200 (2006).

57 Kastner, P., Aukenova, A. & Chan, S. Evolution of the Ikaros family transcription factors: From a deuterostome ancestor to humans. Biochem Biophys Res Commun 694, 149399, doi:10.1016/j.bbrc.2023.149399 (2024).

58 Schmitges, F. W. et al. Histone methylation by PRC2 is inhibited by active chromatin marks. Mol Cell 42, 330–341, doi:10.1016/j.molcel.2011.03.025 (2011).

59 DelRosso, N. et al. Large-scale mapping and mutagenesis of human transcriptional effector domains. Nature 616, 365–372, doi:10.1038/s41586-023-05906-y (2023).

60 Apostolov, A. et al. Sumoylation Inhibits the Growth Suppressive Properties of Ikaros. PLoS One 11, e0157767, doi:10.1371/journal.pone.0157767 (2016).

61 Li, S. et al. ITGA5 Is a Novel Oncogenic Biomarker and Correlates With Tumor Immune Microenvironment in Gliomas. Front Oncol 12, 844144, doi:10.3389/fonc.2022.844144 (2022).

62 Zhu, H., Wang, G., Zhu, H. & Xu, A. ITGA5 is a prognostic biomarker and correlated with immune infiltration in gastrointestinal tumors. BMC Cancer 21, 269, doi:10.1186/s12885-021-07996-1 (2021).

63 Cassinat, B. et al. CCND2 mutations are infrequent events in BCR-ABL1 negative myeloproliferative neoplasm patients. Haematologica 106, 863–864, doi:10.3324/haematol.2020.252643 (2021).

64 Li, Z. et al. GDNF family receptor alpha 2 promotes neuroblastoma cell proliferation by interacting with PTEN. Biochem Biophys Res Commun 510, 339–344, doi:10.1016/j.bbrc.2018.12.169 (2019).

65 Shi, H. et al. BCL11A Is Oncogenic and Predicts Poor Outcomes in Natural Killer/T-Cell Lymphoma. Front Pharmacol 11, 820, doi:10.3389/fphar.2020.00820 (2020).

