## Supplementary Figures for "Conserved helical motifs in the Ikaros IDR mediate NuRD interaction and transcriptional repression"

**Supplementary Figure 1**

a) Volcano plot of proteins identified in anti-HA ChIP-MS of HA-Ikaros expressing B3 cells and untransduced B3 control.
b) Ikaros-ERt2 affinity purification MS interactors performed in absence of crosslinking displayed by abundance, with Ikaros (Ikzf1) in maroon, Aiolos (Ikzf3) in red, NuRD subunits in blue, Kap1 in yellow, FACT subunits in green, and SAGA SWI/SNF subunits in orange. Average of 3 replicates.
c) Volcano plot of proteins identified in anti-HA affinity purification MS of HA-Ikaros-ERt2 expressing B3 cells (with 2h of 4-OHT induction) compared to untransduced B3 control.

**Supplementary Figure 2**

a) Heatmap of differentiatially expressed genes (LRT padj <0.01 and log2FC>1 or <-1) over 24h nascent chromatin-associated RNA-seq timecourse following Ikaros induction performed in duplicate, and K-means clusters of genes by behaviour into C1 Fast Repressed, C2 Slow Repressed, C3 Activated, C4 Slow Activated, and C5 non-monotonic
b) Gene ontology of top pathways enriched in C1 Fast Repressed, C2 Slow Repressed, C3 Activated, C4 Slow Activated and C5 non-monotonic genes.
c) Ikaros motif density (GGAA and GGGA count/bp) at sites with early significant decreased accessibility, increased accessibility, or unchanged accessibility.
d) Homer motif analysis showing top enriched motifs at ATAC NFR sites with early significant increased accessibility compared to regions with unchanged accessibility.
e) Overlap of sites with early significant decreased accessibility, increased accessibility or with unchanged accessibility with H3K27ac peaks and ROSE called superenhancers based on H3K27ac ChIP-seq.
f) Heatmap and metaprofile plot showing chromatin accessibility and H3K27ac at 0hm 30m, 1h, and 2h following Ikaros induction at +/-1kb around sites centered at ATAC peaks with early decreased, increased or unchanged accessibility.
g) Metaprofile plot showing KAP1 binding and H3K9me3 levels at 0h and 1h following Ikaros induction at regions +/-1kb of sites centered at ATAC peaks with early decreased, increased or unchanged accessibility.
h) IGV browser tracks showing KAP1 (purple) and H3K9me3 (pink) at genomic regions with Ikaros (red) and decreased accessibility ATAC (blue), and sites without Ikaros.

**Supplementary Figure 3**

a) Proportion of sites that show early significant decreased or increased chromatin accessibility at active promoters vs enhancers.
b) Proportion of enhancers (dark green) and promoters (light green) at sites showing significant loss of H3K27ac (padj<0.01) and greater than 1.5 fold and 2 fold loss of H3K27ac.
c) Number of Ikaros peaks at enhancers in the region +/- 20kb TSS of Fast Repressed, Slow Repressed, Activated, Slow Activated, and non differentially expressed genes. Pairwise Wilcoxon rank sum test was performed for Fast Repressed vs other gene class with Bonferroni corrected p-adjusted values stated.
d) Proportion of Enhancer – Promoter connections for Fast Repressed, Slow Repressed, Activated, Slow Activated, and non-differentially expressed genes, where AR is the correlation between enhancer and promoter activity and C is a measure of contact frequency between the enhancer and promoter (see Methods)
e) Proportion of enhancer ATAC peaks within +/- 20kb of TSS Fast Repressed, Slow Repressed, Activated, Slow Activated, non-differentially expressed, and randomly sampled genes that show significant (padj<0.01) decreased or increased accessibility within 2h of Ikaros induction.
f) Odds ratio and significance for enhancer accessibility loss or gain of Fast Repressed, Slow Repressed, Activated or Slow Activated genes compared to a randomly sampled set of genes.
g) Odds ratio and significance for enhancer H3K27ac loss or gain of Fast Repressed, Slow Repressed, Activated or Slow Activated genes compared to a randomly sampled set of genes.

**Supplementary Figure 4**

a) Alphafold structure of the Ikaros protein with the conserved helices containing the 9aa motifs in yellow, DNA binding ZnF1-4 in blue, and dimerization ZnF5-6 in green.
b) The hydrophobicity profile for the three Ikaros 9aa motifs sequences versus an ideal prediction.
c) Western blot showing equivalent levels of Ikaros and mutant expression in B3 cells.
d) Schematic of Ikaros mutants ΔED, ΔED1 (Helix1 deletion), and ΔED2 (Helix2 deletion).
e) qPCR of normalised gene expression at several Fast Repressed genes before and after induction with wildtype Ikaros or mutants ΔED, ΔED1, and ΔED2.
f) Quantification of NuRD subunits abundance (normalized to same scaled) affinity purification-MS of full length Ikaros, ΔED, ΔPEST, and untransduced cells.

**Supplementary Figure 5**

a) Protein sequence alignment of the “KRKSSMPQ” motif present in Ikaros family TFs (Ikzf1, Ikzf2, Ikzf3, Ikzf4) with the 12 aa motif identified in NuRD-interacting TFs Fog1, Bcl11a, Sall4.
b) The 12 aa motif is formed of 3 positively charged residues (KRK or RRK) followed by a largely polar stretch of residues
c) AlphaPullDown prediction of Ikaros 235-362 (conserved helical motif region) with the NuRD subunit RBBP4. Interface pTM score of 0.73.
d) Comparison of the predicted structure of Ikaros-RBBP4 (with obstructing Helix1 residues hidden) with the crystal structures of Fog1 and Bcl11a N-terminal peptides with RBBP4.
e) Close up of the KRK/RRK interaction with the central pocket of RBBP4.

#### Supplementary Figure 1

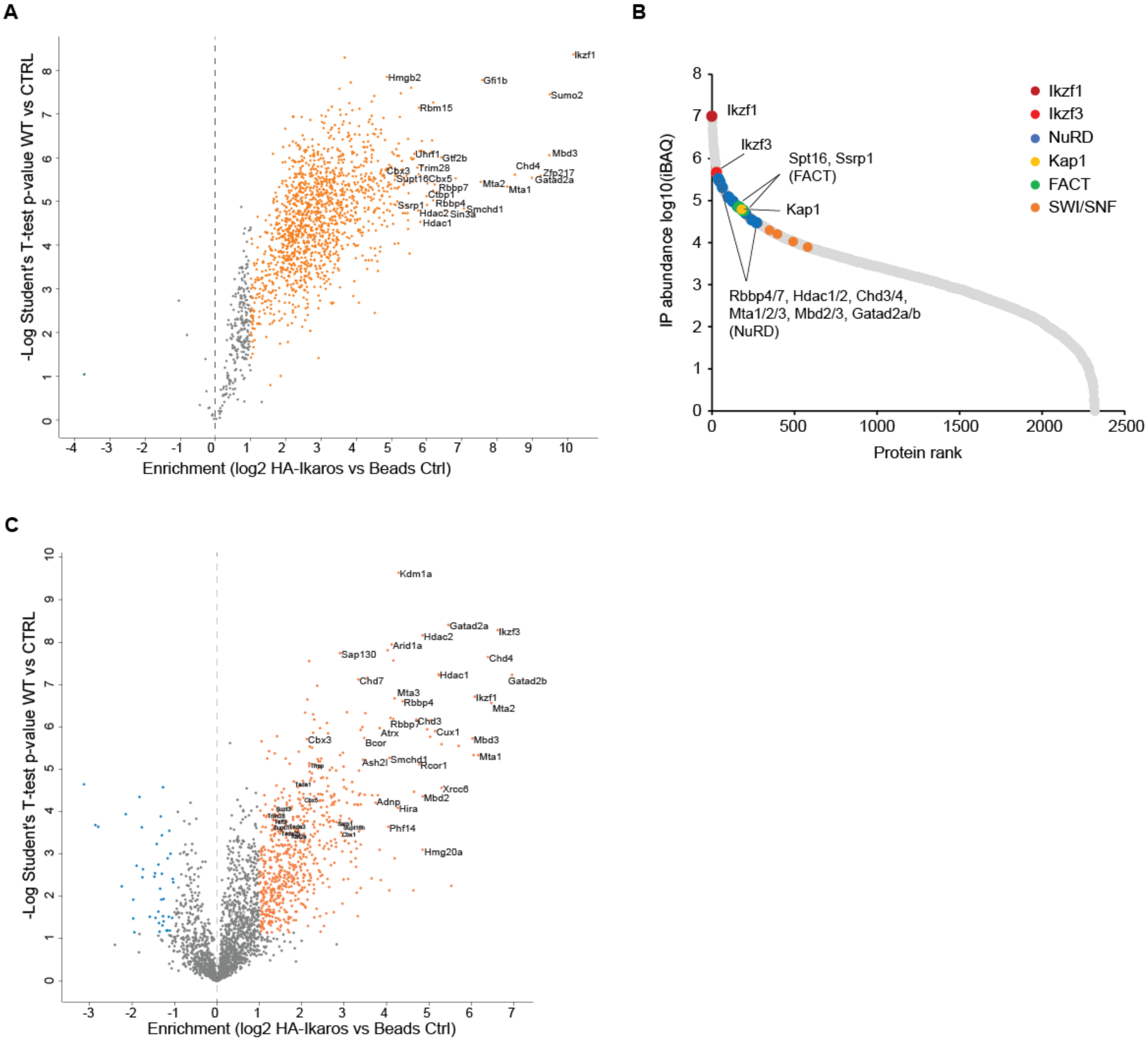

### Supplementary Figure 2

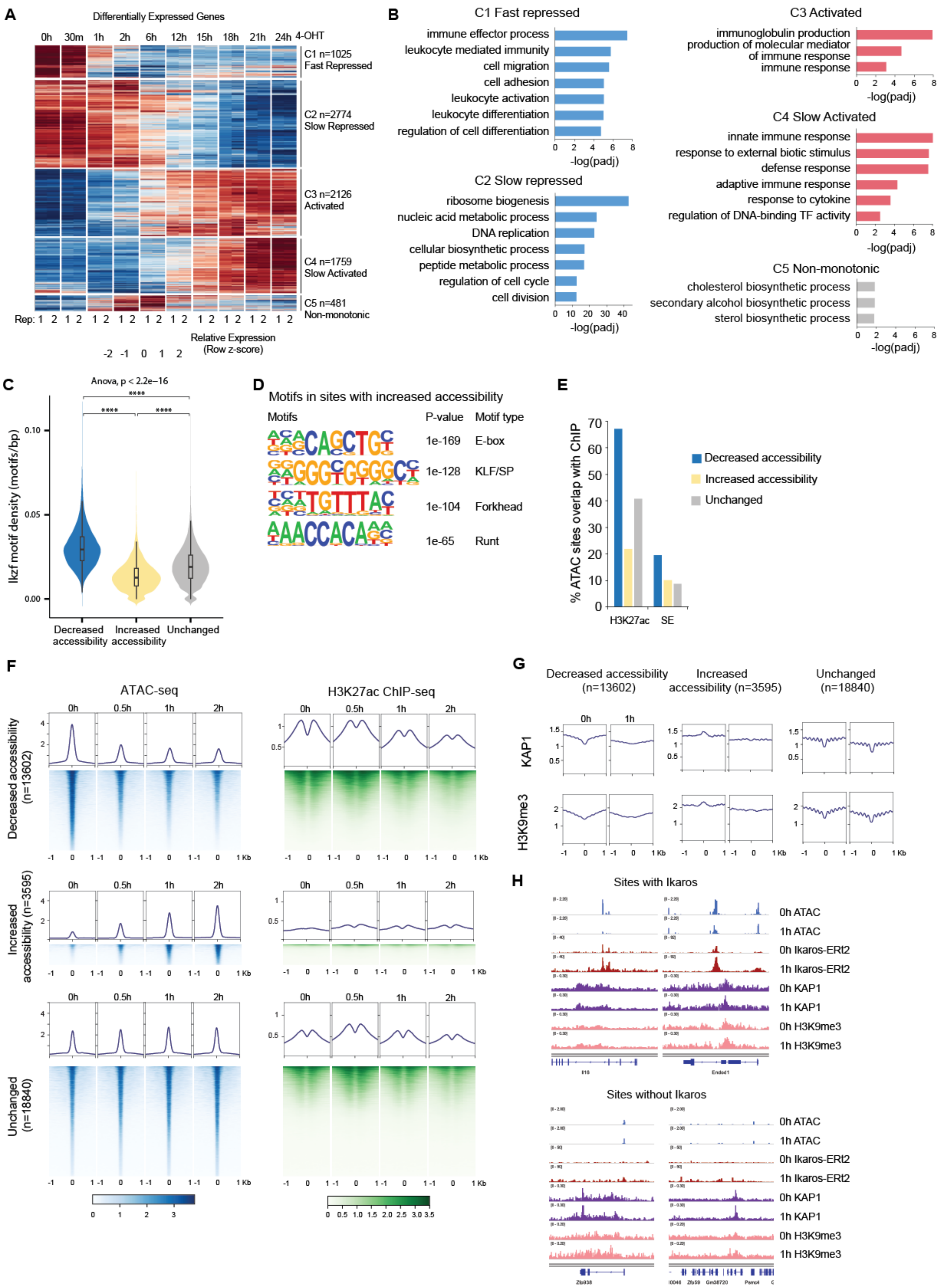

### Supplementary Figure 3.

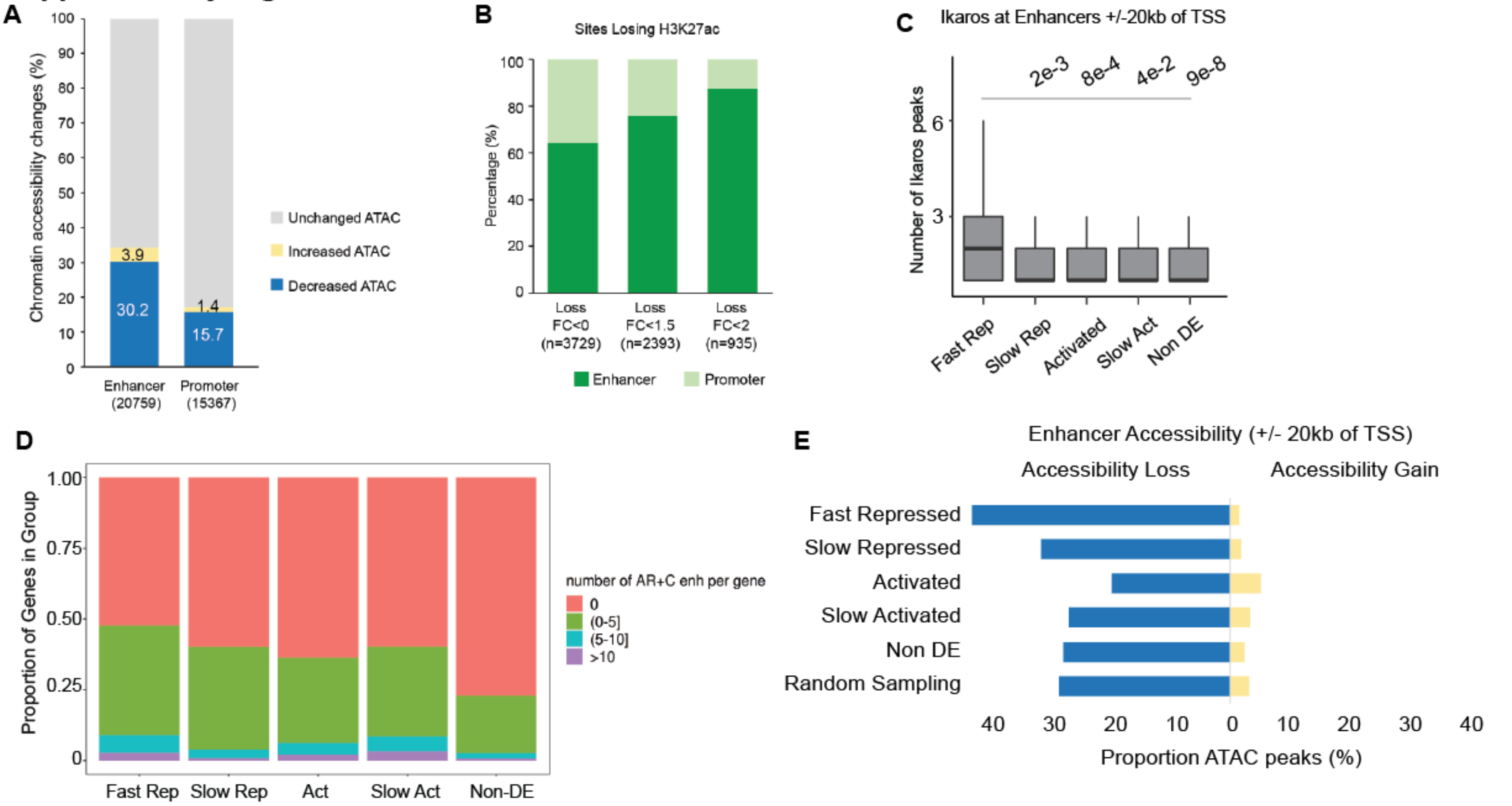


**F**

| Enhancer | Accessibility loss | | Accessibility gain | |
|---|---|---|---|---|
| | Odds Ratio | Padj. | Odds Ratio | Padj. |
| Fast Repressed | 1.90 | <0.0001 | 0.47 | 0.01 |
| Slow Repressed | 1.16 | n.s. | 0.58 | n.s. |
| Activated | 0.61 | <0.0001 | 1.68 | 0.03 |
| Slow Activated | 0.92 | n.s. | 1.06 | n.s. |

**G**

| Enhancer | H3K27ac loss | | H3K27ac gain | |
|---|---|---|---|---|
| | Odds Ratio | Padj. | Odds Ratio | Padj. |
| Fast Repressed | 2.94 | <0.0001 | 0.25 | 0.0002 |
| Slow Repressed | 1.12 | n.s. | 0.66 | n.s. |
| Activated | 0.37 | <0.0001 | 3.60 | <0.0001 |
| Slow Activated | 1.07 | n.s. | 1.57 | 0.0243 |

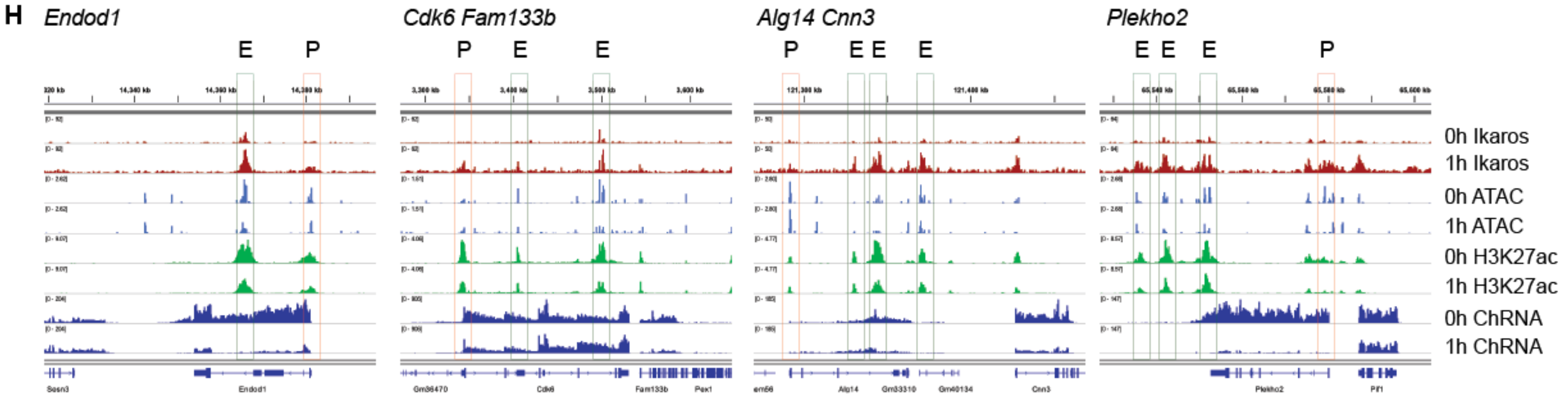

**Supplementary Figure 4.**

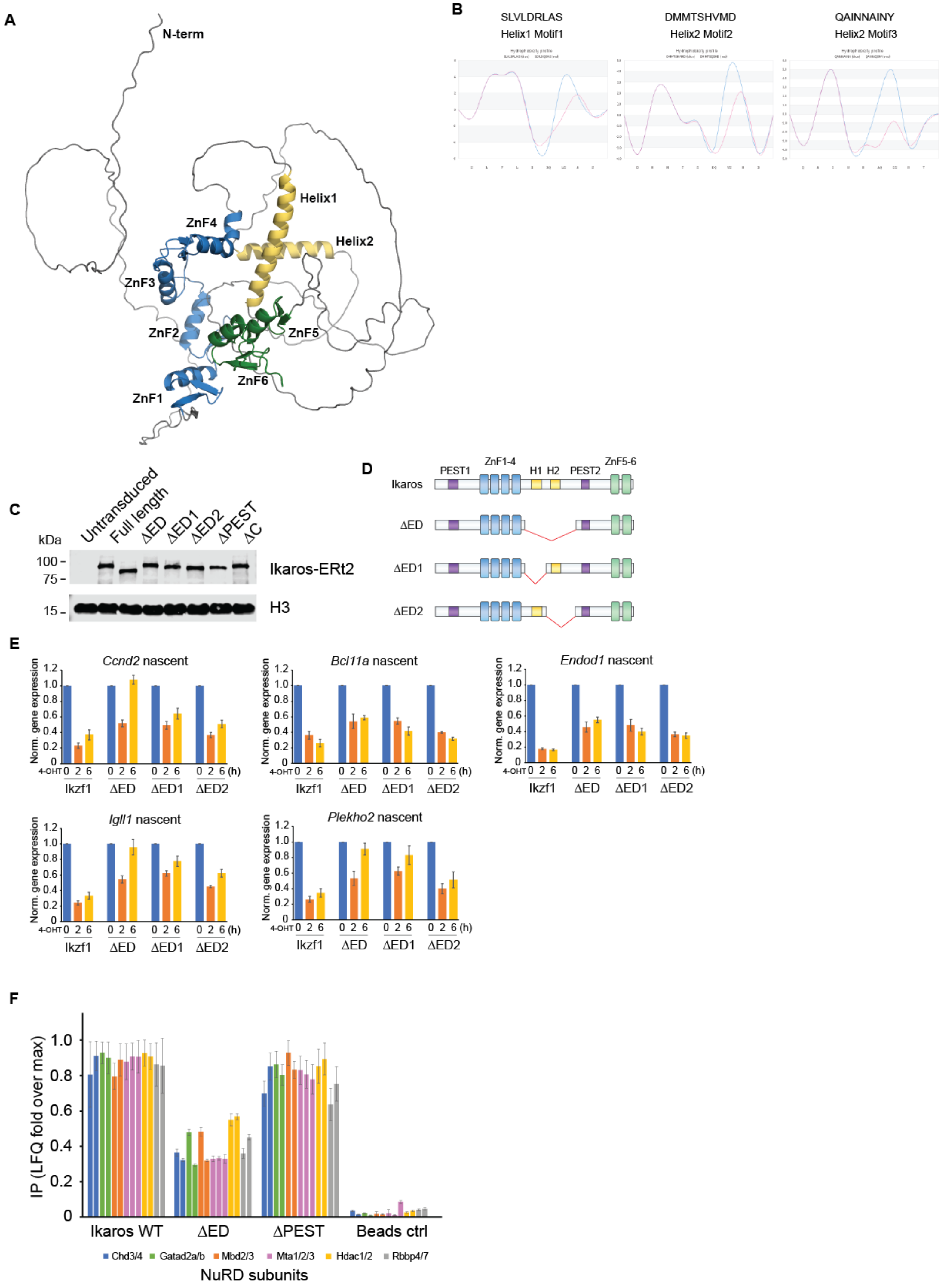

#### Supplementary Figure 5

**A**

```
VAKRKSSMPQKF   Ikzf1
VAKRKSSMPQRF   Ikzf1 (zebrafish)
VAKRKSSMPQKF   Ikzf3
LTKRKRSTPQKF   Ikzf4
MGKRKSSTPQKF   Ikzf2
MSRRKQSNPRQI   Fog1
MSRRKQGKPQHL   Bcl11a
MSRRKQAKPQHI   Sall4
: :** . *:::
```

**B**

| | | positive | | | polar | | | | polar | | | |
|---|---|---|---|---|---|---|---|---|---|---|---|---|
| - | A | R | T | K | Q | T | A | R | K | S | T | H3 |
| V | A | K | R | K | S | S | M | P | Q | K | F | Ikzf1 (Ikaros) |
| V | A | K | R | K | S | S | M | P | Q | K | F | Ikzf3 (Aiolos) |
| M | S | R | R | K | Q | S | N | P | R | Q | I | Fog1 |
| M | S | R | R | K | Q | G | K | P | Q | H | L | Bcl11a |
| M | S | R | R | K | Q | A | K | P | Q | H | I | Sall4 |

**C**

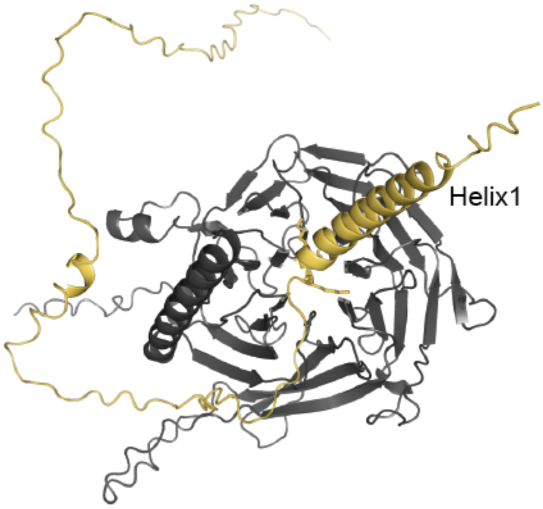


**D**

Ikaros KRK - RBBP4 model (Helix1 hidden) | Fog1 RRK - RBBP4 (PDB 2xu7) | Bcl11a RRK - RBBP4 (PDB 5vtb)

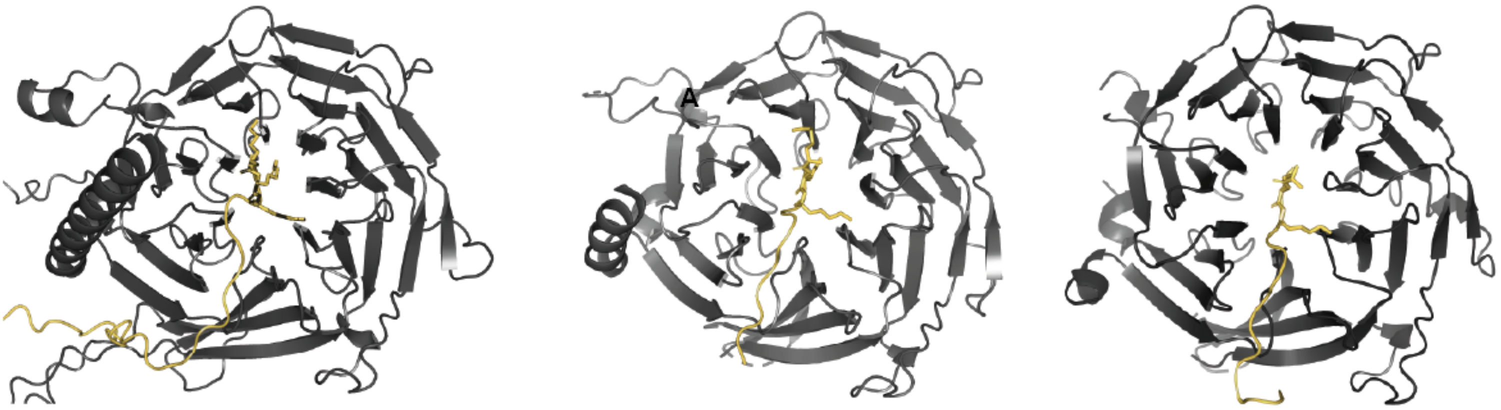


**E**

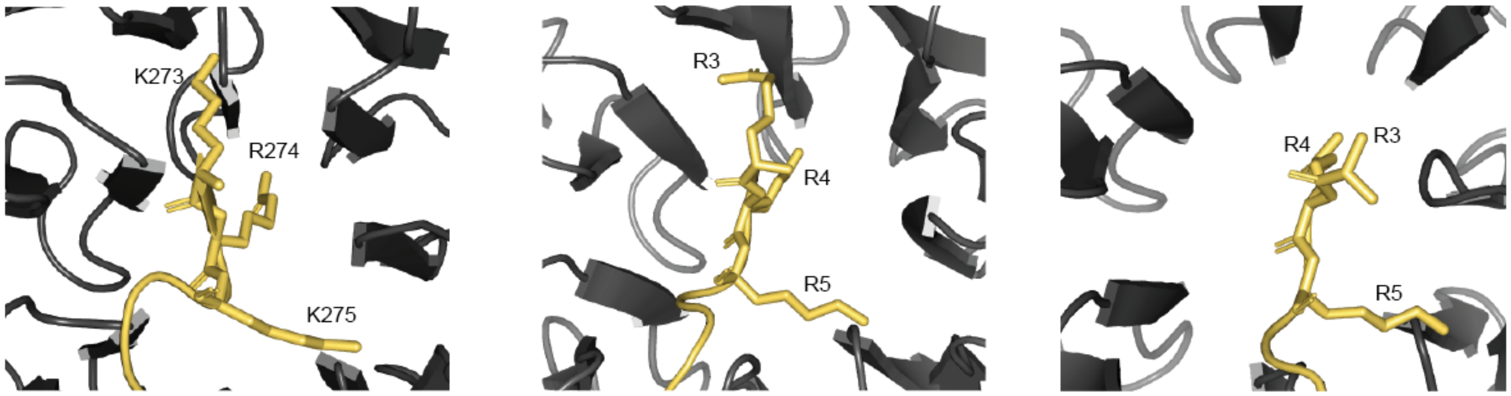
