## Supplementary material for "Conserved helical motifs in the Ikaros IDR mediate NuRD interaction and transcriptional repression": Materials and Methods

#### Cell culture

B3 pre-B cells were maintained in IMDM with 10% heat inactivated FBS and 1x Pen/Strep (Gibco™ 15070063) and 1x L-glutamine (Gibco™ 25030081) between 0.5 – 2 million/ml. HEK293T cells were maintained in DMEM with 10% heat inactivated FBS and 1x Pen/Strep and 1x L-glutamine, and split every 2-3 days. B-ALL cell lines BV173 and SupB-15 were grown in RPMI with 10% heat inactivated FBS and 1x Pen/Strep and 1x L-glutamine and maintained between 0.2 – 1.5 million/ml. All cells were frozen in 90% heat inactivated FBS with 10% DMSO and stored long term in LN2.

#### Retrovirus and Lentivirus infection

The inducible Ikaros-ERt2 construct was introduced in the B3 cell line using retroviral transduction. HEK293T at 80% confluency in 10 cm dish were transfected with 4μg of the transfer plasmid MSCV-Ikaros-ERt2-IRES-GFP and 4μg of pCL-Eco retroviral packing plasmid using the calcium phosphate method. Viral supernatant was collected every 12h from 24h to 72h after transfection, pooled and filtered in a low protein bind 0.45 μm filter (Millipore SLHV033N). For retroviral spinfection, 3ml of viral supernatant was added to 2 million B3 cells with 4 μg/ml polybrene (TR-1003-G) and spinfection was performed at 2500rpm for 1.5h at 30 °C. B3 cells were sorted for GFP 3-4 days post infection. 500nM of 4-hydroxytamoxifen (Sigma-Aldrich H7904-5MG) was used to induce nuclear translocation of Ikaros.

The inducible Ikaros-ERt2 construct was introduced in the B-ALL cell lines BV173 and SupB15 using lentiviral transduction. Low passage Lenti-X HEK293T cells at 80% confluency seeded in 15 cm dishes were transfected with 20 μg transfer plasmid pLVX-Ikaros-ERt2-IRES-ZsGreen and 18 μg of psPAX2 packaging vector and 7 μg pMD2.G envelope vector using Lipofectamine3000. Collection and spinfection were every 12h from 24h to 72h after transfection, pooled, filtered in a low protein bind 0.45 μm filter, and concentrated 3 times using a 100 kda MWCO spin concentrator. Lentiviral spinfection was performed with 3ml of concentrated viral supernatant was added to 2 million BV173 or SupB15 cells with 4 μg/ml polybrene (TR-1003-G) and spinfection was performed at 2500rpm for 1.5h at 30 °C. Cells were sorted for ZsGreen 4 days post infection. 500nM of 4-hydroxytamoxifen (Sigma-Aldrich H7904-5MG) was used to induce nuclear translocation of Ikaros.

#### Cell growth assay

B-ALL were seeded at 0.2 million/ml into 12-well plates in a total volume of 1ml in cell culture media (RPMI with 10% heat inactivated FBS and 1x Pen/Strep and 1x L-glutamine) supplemented with 500 nM 4-OH tamoxifen. Cells were counted every other day using the Countess Cell Counter after resuspension.

**Chromatin-associated RNA-seq sample preparation**

Subcellular fractionation and isolation of chromatin bound nascent RNAs was adapted from Nesterova et al 2019[1]. Briefly, cell nuclei were isolated in lysis buffer with a sucrose cushion. Nuclei were then lysed in a urea buffer, and the insoluble pellet containing the chromatin fraction and associated nascent RNAs was purified using the Zymo Direct-zol RNA Miniprep Kit (R2051). Library prep was performed using the NEBNext® Ultra™ II Directional RNA Library Prep Kit for Illumina® (E7760S) using the rRNA depletion (E6310L) workflow and 75bp paired end sequencing was performed on the Ilumina Nextseq 500.

**Chromatin-associated RNA-seq analysis**

Paired-end 75bp sequencing reads were aligned against mouse genome (GRCm38) with nextflow nfcore/rnaseq:1.3 using star as aligner. Gene based read counts were obtained using featureCounts function from Rsubread Bioconductor package and intronic reads were also included in the gene level quantification. Normalisation was performed in DESeq2 Bioconductor package and data was rlog transformed to allow for visualisation by PCA and heatmaps. Differential gene expression (DEG) analysis was conducted using DESeq2 with the DESeq function with likelihood ratio test using parameter test= "LRT". DEGs were defined as those with Benjamini-Hochberg adjusted p-value (P) < 0.01 and a fold change >2 or <0.5 at any of the time points when compared to 0 hours. Heatmaps were generated using R Bioconductor package ComplexHeatmap (Gu et al., 2016[2]). We created five k-means DEG clusters with the parameter row_km = 5 with Heatmap function with set.seed(123) activated in R. Gene ontology analysis for each gene clusters were performed with goseq Bioconductor package (1.24; Young et al., 2010[3]).

**ATAC-seq sample preparation**

ATAC-seq was performed based on the omni-ATAC protocol. Briefly, for each sample 50 000 cells were lysed (10mM Tris pH 7.5, 10 mM NaCl, 3 mM $MgCl_2$, 0.1% NP-40, 0.1% Tween, 0.01% Digitonin), and nuclei were tagmented with 3 ul of Ilumina Tn5 enzyme (Illumina, catalog # 15027865) in TD Buffer with 0.01% Tween and 0.01% Digitonin for 30 minutes at 37°C. DNA was purified using the Zymo DNA Clean and Concentrator and libraries were generated with 7 cycles of PCR with index primers described in the original Buenrostro et al. ATAC-seq protocol. Clean-up was performed with Ampure beads and quantification and quality control of libraries was performed using DNA Qubit and HS DNA Bioanalyser. 75bp Pair-end sequencing was performed on the Ilumina Nextseq 500.

**ATAC-seq analysis**

Paired-end 76 bp ATAC-seq sequencing were proceeded using RTA version 2.4.11 under NextSeq500, with default filter and quality settings for basecalling. The reads were demultiplexed with bcl2fastq 2.20.0, allowing 0 mismatches. After demultiplexing, raw reads were processed with Nextflow nf-core/atacseq:1.0.0 (as in

Ewels et al [4]) with parameters: '--genome GRCm38 -profile conda' using bwa (v0.7.17; Li and Durbin, 2009 [5]) aligner. Processed BAM files from this pipeline was taken forward to perform downstream analysis. ATACshift mode of deeptools (v. 2.3.5; Ramírez et al., 2016 [6]) alignmentSieve function was used to shift the reads with parameters: '--minFragmentLength 0 --maxFragmentLength 120' for Nucleosome Free Region (NFR – fragment length less than 120 bp). The filtered bam files were further processed with deeptools alignment Sieve function with parameters: '--ATACshift --minFragmentLength 180' for Nucleosome Bound Region (NBR – fragment length more than 180 bp). We used the NFR bin for the downstream analysis. Peak calling was proceeded with MACS2 (v2.1.2) callpeak function with parameters: '-f BAMPE -g mm' (Zhang et al., 2008 [7]). Consensus peaks throughout different time points were defined by using nf-core/atacseq. Peaks overlapped with the black list mm10 v1 (Amemiya et al., 2019 [8]; ENCODE Project Consortium, 2012[9]) were removed from downstream analysis. Peak-based read counts were then obtained using featureCounts (v1.6.4; Liao et al., 2014[10]). Differential accessibility analysis was proceeded with DESeq2 (Love et al., 2014[11]) Bioconductor package. Peaks were annotated with ChIPseeker Bioconductor package (Yu et al., 2015[12]).

BigWig files were generated based on BAM files from the nucleosome free regions (NFR). Scale factors for each BAM file were calculated beforehand using the formula 1000000 divided by the number of mapped reads obtained from the Samtools (v.1.9; Danecek et al., 2021[13]) flagstat function. genomeCoverageBed function from Bedtools (v2.27.1; Quinlan and Hall, 2010[14]) was applied with command: 'genomeCoverageBed -ibam $outDir/${SAMPLE}.mLb.ss.sorted.NFR.bam -bg -scale $SCALE_FACTOR -pc | sort -k1,1 -k2,2n > $outDir/bigwig_gcov/${SAMPLE}.mLb.ss.sorted.NFR.bedGraph'. Followed by deploying bedGraphToBigWig (Kent et al., 2010) with the following command: 'bedGraphToBigWig $outDir/bigwig_gcov/${SAMPLE}.mLb.ss.sorted.NFR.bedGraph $gsizes $outDir/bigwig_gcov/${SAMPLE}.mLb.ss.sorted.NFR.bigWig'.

**Chip-seq sample preparation**

ChIP-seq was adapted from Zhang et al [15]. For calibrated ChIP-seq 2 million human HEK293T cells were mixed with 45 million B3 cells prior to fixation as in Fursova et al[16]. Cells were fixed at room temperature in 10ml PBS with 1mM DSG (ThermoFisher 20593) for 30 min followed by addition of 1% formaldehyde (ThermoFisher 28908) and incubated for a further 10min. The reaction was quenched with 125mM glycine. Cells were lysed with LB1 (50mM Hepes pH 7.5, 140 mM NaCl, 1mM EDTA, 10% glycerol, 0.5% NP40, 0.25 Triton X-100, fresh protease inhibitor). Nuclei was pelleted and washed once in LB2 (10mM Tris pH 8.0, 200mM NaCl, 1mM EDTA, 0.5mM EGTA, fresh protease inhibitor) then resuspended in LB3 (10 mM Tris pH 8.0, 100mM NaCl, 1mM EDTA, 0.5mM EGTA, 0.1% sodium deoxycholate, 0.5% N-lauroylsarcosine or without detergent when MNase was used). Chromatin was digested by sonication or MNase so that majority of fragments were between 100-400bp. Chromatin equivalent to 4 million cells was used per IP. Chromatin samples were diluted in ChIP DB (20mM Tris pH 8, 150mM NaCl, 1% Triton x100, 1mM EDTA, fresh protease inhibitor) and a small Input sample was reserved. HA-Ikaros-ERt2 ChIP was performed using overnight incubation with Pierce magnetic anti-HA beads (Pierce 88836). 5 $\mu$g of antibodies for H3K27ac

(Active Motif 39133) and CHD4 (Abcam ab70469), H3K9me3 (Diagenode C15410193), and KAP1 (Abcam ab10483) were incubated with each reaction overnight, followed by 2h of incubation with magnetic Prot A/G Dynabeads (ThermoFisher 10001D, 10003D). Washes were performed for 5 min rotating in the cold room with each buffer, Low Salt Buffer (0.1% SDS, 1% TritonX-100, 2mM EDTA, 20mM Tris pH 8, 150mM NaCl), High Salt Buffer (0.1% SDS, 1% TritonX-100, 2mM EDTA, 20mM Tris pH 8, 500mM NaCl), LiCl Buffer (250mM LiCl, 1% NP-40, 1% sodium deoxycholate, 1mM EDTA, 10mM Tris pH 8) and TE (10mM Tris pH 8, 1mM EDTA). DNA was eluted with 1% SDS and 0.1M $NaHCO_3$. For reverse crosslink, NaCl was added to a final concentration of 0.2M, and samples were incubated at 65°C shaking overnight, followed by 2h of incubation with 200μg/ml RNaseA then 2h with 200μg/ml Proteinase K treatment at 45°C. DNA was eluted off the beads with fresh EB (1% SDS and 0.1M $NaHCO_3$) then purified using the Zymo ChIP DNA Clean and Concentrator (Zymo D4013). NEBNext® Ultra™ II DNA Library Prep Kit for Illumina® (E7645S) was used for all ChIP-seq libraries with PCR amplification of 5-8 cycles. Clean-up was performed with Ampure XP beads (Beckman A63882) and quantification and quality control of libraries was performed using DNA Qubit and HS DNA Bioanalyser. 40bp paired end sequencing was performed on the Ilumina Nextseq 500.

All NGS sequencing reads in this study were proceeded using RTA version 2.4.11 on the NextSeq500, with default filter and quality settings for basecalling. The reads were demultiplexed with bcl2fastq 2.20.0, allowing 0 mismatches.

**ChIP-seq analysis**

*Ikaros, H3K27ac, and CHD4 0, 1, and 2 hours* Raw paired-end 41bp sequencing reads were aligned against mouse genome (mm10) using Bowtie2 (v. 2.4.2; Langmead and Salzberg, 2012[17]) with parameters: '--no-mixed --no-discordant'. Picard (v. 2.23.7; https://broadinstitute.github.io/picard/) SortSam and MarkDuplicates were applied for sorting and converting sam files to bam files and duplicated reads were marked. Peak calling was proceeded with MACS2 (v.2.2.7.1) callpeak function (with -f BAMPE -g mm for Ikaros and H3K27ac; with -f BAMPE --broad -g mm for CHD4). If a peak appears in 1 sample at any of the timepoints in the datasets, we treat it as a valid peak. The consensus peaks across different time points were defined by using the Bioconductor package GenomicRanges (Lawrence et al., 2013[18]) with the reduce function. Peak-based read counts were then obtained using the featureCounts function from Rsubread (Liao et al., 2019[10]) Bioconductor package. Differential binding analysis was proceeded with DESeq2 Bioconductor package. For generation of Bigwig files, we used the IP raw concentration at 2 hours from each ChIPs as the baseline to calculate the size factor (SF) for the downsampling analysis at each time point. The SF at each time point was calculated by dividing the IP raw concentration at time 0 hours or 1 hour by the IP raw concentration at 2 hours. Reads from raw fastq files were randomly selected based on the SF calculated for downsampling by using seqtk (v1.3; https://github.com/lh3/seqtk) with 'seqtk sample -s100 $ori.fastq.gz $sizeFactor - | gzip - > $DS_fastq.gz'. After downsampling the fastq files, we applied the same analytical pipeline described in this section for alignment, sorting, converting to BAM files, and marking duplicated reads. We generated downsampled BigWig files using Deeptools (v. 3.5.0) with the following command: 'bamCoverage -b ChIP.DupMark.bam -p 16 -bl mm10-blacklist_v1.bed --extendReads --effectiveGenomeSize 2652783500 -o ChIP_MarkedDup.bw'.

For calibrated ChIP-seq samples KAP1 and H3K9me3, we analyzed this human-calibrated ChIP dataset based on the strategy by Fursova *et al.* (2019)[16]. In summary, raw paired-end 41bp sequencing reads were aligned to a merged genome of mouse (mm10) and human (hg38) using Bowtie2 (v. 2.4.2), with prefixes Mm10 and Hg38 for mouse and human chromosome names, respectively. We employed Picard (v. 2.23.7) SortSam and MarkDuplicates to sort and convert the SAM file to BAM files, marking duplicated reads. The Samtools view function was applied to separate human and mouse-mapped reads using the following commands: samtools view -h -L $bedfile_hg38Chrs or samtools view -h -L $bedfile_mm10Chrs. We removed reads that mapped to both human and mouse and then separated the remaining reads based on whether they mapped to human or mouse. The downsampling factors (DFs) can be calculated based on this equation (Fursova et al., 2019[16]):

$$DF = \alpha \times \frac{1}{N(spikeIn\ in\ ChIP)} \times \frac{N(\text{spikeIn in Input})}{N(\text{mouse in Input})}$$

N (spikeIn in ChIP) - total number of reads aligning to the spike-in genome for each ChIP-seq sample;

N (spikeIn in Input) - total number of reads aligning to the spike-in genome in the corresponding Input sample;

N (mouse in Input) - total number of reads aligning to the mouse genome in the corresponding Input sample;

α - coefficient applied for all the files normalized together so the value of the largest downsampling factor equals 1.

Normalised bigwig files for ChIPs were generated based on the DFs calculated by the equation above using Deeptools (v.3.5.0) bamCoverage function with parameters -bl $blacklist --binSize 1 --extendReads --effectiveGenomeSize 2652783500 --scaleFactor $showDF.

**Published NGS data**

Bed files containing peak information for the following published NGS experiments were directly retrieved from GEO. Peaks that overlap with the blacklist were removed from the subsequent analysis.

| Dataset | genome | GEO accession | Reference | Source |
|---|---|---|---|---|
| Ctcf ChIP | mm10 | GSE109671 | Koohy et al., 2018[19] | Pre-B cells from young mice |

| | | | | |
|---|---|---|---|---|
| Pre-B domains | mm9 | GSE63525 | Rao et al, 2014[20] | Pre-b lymphoblast CH12-LX |

### Super Enhancer identification

H3K27ac peaks were utilized in the identification of super enhancers through the Rank Ordering of Super-Enhancers (ROSE; v.1.0.0) method (https://github.com/stjude/ROSE; Lovén *et al.*, 2013[21]; Whyte *et al.*, 2013[22]) with parameters -s 12500 -t 0.

### Enhancer Promoter connectivity analysis

Activation ratio + contact (AR + C) is a computational method for enhancer-promoter (E-P) assignment based on two components: 1) Activation ratio which captures how well the enhancer activity predicts the gene promoter activity across a panel of tissues / cell types and 2) Contact: by including information about the frequency of 3D contacts between the enhancer and promoter.

Activation Ratio: Input data for AR calculation was based on enhancer and promoter FANTOM5 CAGE data[23] across 395 mouse primary cell, cell line and tissue samples. To identify enhancers that are predictive of gene promoter activity, the level of co-expression between putative enhancers and all promoters within a linear distance of 3 Mb was quantified and statistically evaluated. This done by applying the following procedure to each E-P pair: 1) The enhancer was annotated as "active" or "inactive" in each human individual biosample constituting the tissue / cell type panel. More specifically, the enhancer was labelled as "inactive" if its CAGE expression level was equal to zero TPM, and "active" otherwise. 2)The median CAGE expression level (in TPM) of the candidate target promoter was recorded for samples in which the enhancer was active (E+) and samples in which the enhancer was inactive (E-). Next, the activation ratio (AR) for that E-P pair was defined as the log-fold change in median promoter expression in E+ samples compared to E- samples. 3) The statistical significance of the observed activation ratio for that E-P pair was evaluated based on a permutation test. This was done by selecting a random set of E+ samples of the same size as the original E+ set, annotating all remaining samples as E-, and re-estimating the activation ratio. This was repeated 10,000 times and the fraction of permuted activation ratio values that are at least as extreme as the observed activation ratio (absolute value) was estimated and used as an empirical P-value. 4) Empirical P-values are then subjected to multiple testing correction. To do this, a 10% FDR (Benjamini-Hochberg) correction was applied to all tests performed genome-wide, equal to the total number of E-P pairs tested. E-P pairs below the significance threshold are considered as significantly co-expressed.

Contact: Enhancer-promoter pairs supported by 3D contact were identified using high-resolution Hi-C or Micro-C data from different cell types to identify enriched contacts between pairs of genomic bins and subsequently overlapping those with enhancers and promoters from the previous section. More specifically, we used Hi-C matrices from mESCs and neural progenitor cells[24], a Hi-C matrix from pre-B cells[25], and a Micro-C matrix from JM8 NP4 cells[26]. An individual E-P pair was marked as "supported by 3D contact" in each of the 4 cell types if its normalised contact frequency exceeded a predefined threshold representing the genome background contact frequency in that specific cell type (or contact matrix). All E-P pairs that were supported by 3D contact in at least one of the 4 cell types were kept for further analysis.

Combining AR + C: The final set of AR+C associations was defined by taking all E-P pairs that had a significant positive activation ratio and were also supported by 3D contact data. To obtain a single measure of enhancer responsiveness per gene, the median activation ratio across all its AR+C associations was estimated for each gene. In addition, the number of AR+C enhancers (enhancers for which both the activation ratio and contact criteria are satisfied) was estimated per gene.

**ChIP-MS and Affinity Purification MS sample processing**

ChIP-MS for HA-Ikaros was performed similar to the RIME protocol[27]. Three replicates of 40 million B3 cells, and three replicates of 40 million B3 cells expressed HA-Ikaros were fixed in 1% formaldehyde in PBS for 8 minutes rotating at room temperature, then quenched with 125mM glycine. Cells were lysed with LB1 (50mM Hepes pH 7.5, 140 mM NaCl, 1mM EDTA, 10% glycerol, 0.5% NP40, 0.25 Triton X-100, fresh protease inhibitor). Nuclei was pelleted and washed once in LB2 (10mM Tris pH 8.0, 200mM NaCl, 1mM EDTA, 0.5mM EGTA, fresh protease inhibitor) then resuspended in LB3 (10 mM Tris pH 8.0, 100mM NaCl, 1mM EDTA, 0.5mM EGTA, 0.1% sodium deoxycholate, 0.5% N-lauroylsarcosine) sonicated for 15 min 30s on 30s off to isolate chromatin fragments in the range of 100-300bp. Nuclear lysate was incubated with Pierce Magnetic anti-HA beads for 5h, and washed a total of 10 times for 3 minutes rotating in the following buffers, three times in Low Salt Buffer (0.1% SDS, 1% TritonX-100, 2mM EDTA, 20mM Tris pH 8, 150mM NaCl), three times in High Salt Buffer (0.1% SDS, 1% TritonX-100, 2mM EDTA, 20mM Tris pH 8, 500mM NaCl), twice in LiCl Buffer (250mM LiCl, 1% NP-40, 1% sodium deoxycholate, 1mM EDTA, 10mM Tris pH 8), and twice in AMBIC buffer prior to downstream processing for proteomic analysis.

AP-MS of inducible wildtype Ikaros and ∆ED and ∆PEST mutants were performed in the absence of fixation. Four replicates were performed for wildtype Ikaros and both mutants, along with control untransduced cells. B3 cells were induced with 500nM of 4-hydroxytamoxifen for 2h (Sigma-Aldrich H7904-5MG) to induce nuclear translocation of Ikaros. The cell pellet was lysed (10mM Tris pH7.5, 10 mM KCl, 1.5 mM $MgCl_2$, 0.1% NP40, 0.1% Tween20, fresh protease inhibitors) then underlaid with lysis buffer + 24% sucrose cushion. Nuclei were pelleted and nuclear proteins were obtained using a standard high salt extraction (10mM Hepes pH 7.5, 5% glycerol, 1.5mM $MgCl_2$, 0.2mM EDTA, 300mM NaCl) by incubating for 1h on ice. Nuclear extracts were treated with benzonase to remove contaminant nucleic acids. For the IP extract was incubated for 5h rotating in the cold room with Pierce Magnetic HA beads, and washed for a total of 10 times, four times in wash buffer (20mM Tris pH 8, 5% glycerol, 200mM NaCl, 0.2% NP-40, fresh protease inhibitors), four times in wash buffer with no detergent, and 2x in 20mM EPPS solution.

For each experiment, 10% of the beads were taken for silver staining, and the remaining for downstream proteomic analysis. A modified on-beads digestion protocol based on Turriziani et al., 2014 [28] was followed for the ChIP-MS samples. The beads were subjected to trypsin digestion in 60µL of buffer 1 (2M urea, 20mM HEPES at pH8 and 1µg trypsin) (Trypsin Gold, Promega, V5280) for 30 minutes at 27°C in a thermomixer, with shaking at 800 rpm. After centrifugation for 1 minute at 7000 rpm, the supernatant was collected into fresh low-bind tubes. Beads were washed twice with 30µL of buffer 2 (2M urea, 20mM HEPES at pH 8 in 1mM DTT), and the supernatants were combined. Overnight digestion was performed at 37°C. The following morning, 5uL of Chloroacetamide (200mM) were added, followed by a 30-minute incubation in the dark. To stop the digestion and facilitate protein digest clean-up, 20µL of 1% trifluoroacetic acid (TFA) were added, to a final concentration 0.1% TFA. Protein digests were de-salted using Glygen C18 spin tips (Glygen Corp, TT2C18.96), eluted with 60% acetonitrile, 0.1% formic acid (FA) with the eluents dried via vacuum centrifugation.

The AP-MS samples were subjected to a non-denaturing on-bead digestion protocol. The beads were washed with 20 mM EPPS pH 8.5 buffer. Briefly, 30 µL of digestion solution containing trypsin (Pierce™ P/N 90059) (20 ng/µL) and LysC (WAKO P/N 129-02541) (10 ng/µL) in 20 mM EPPS pH 8.5 were added to each sample. The samples were placed on a Thermomixer for a 2-hour incubation at 37°C, with shaking at 1100 rpm. After the initial incubation, the samples were placed on a magnetic rack, enabling the separation of the supernatant from the beads, which was then transferred to a fresh PCR plate, for overnight digestion at room temperature.

**Liquid chromatography-tandem mass spectrometry analysis**

ChIP-MS. Chromatographic separation was performed using an Ultimate 3000 RSLC nano liquid chromatography system (Thermo Scientific) coupled to a Thermo Q-Exactive HFX mass spectrometer via an EASY-Spray source. Peptide solutions were injected and loaded onto a trapping column (Acclaim PepMap 100 C18, 100µm × 2cm) for desalting and concentration at 8µL/min in 2% acetonitrile, 0.1% TFA. Peptides were then eluted on-line to an analytical column (EASY-Spray PepMap RSLC C18, 75µm × 50cm) at a flow rate of 250nL/min. Peptides were separated using a 90 minute stepped gradient, 1-22% of buffer B for 60 minutes followed by 22-42% buffer B for another 30 minutes (composition of buffer A – 95/5%: $H_2O$/DMSO + 0.1% FA, buffer B – 75/20/5% MeCN/$H_2O$/DMSO + 0.1% FA) and subsequent column conditioning and equilibration. Eluted peptides were analysed by the mass spectrometer operating in positive polarity using a data-dependent acquisition mode. Ions for fragmentation were determined from an initial MS1 survey scan at 120,000 resolution, followed by HCD (Higher-energy Collision Induced Dissociation) of the top 30 most abundant ions at 15,000 resolution. MS1 and MS2 scan AGC targets were set to 3e6 and 5e4 for maximum injection times of 25ms and 50ms respectively. A survey scan m/z range of 350 – 1750 was used, with normalised collision energy set to 27%, charge state screening enabled with unassigned and +1 charge states rejected. A minimum AGC target was 8e3 with dynamic exclusion set to 50 seconds.

AP-MS. Samples were injected and data acquired in single replicate injections as follows: Chromatographic separation was performed using an Ultimate 3000 RSLC nano liquid chromatography system (Thermo Scientific) coupled to an Orbitrap HFX mass spectrometer (Thermo Scientific) via an EASY-Spray source. Electro-spray nebulisation achieved by interfacing to Bruker PepSep emitters (PN: PSFSELJ20, 20µm). Peptide solutions were injected directly onto the analytical column (self-packed column, CSH C18 1.7µm beads, 300µm × 35cm) at working flow rate of 5µL/min for 8 minutes. Peptides were then separated using a 66 minute stepped gradient: 0-45% of buffer B for 66 minutes (composition of buffer A – 95/5%: $H_2O$/DMSO + 0.1% FA, buffer B – 75/20/5% MeCN/$H_2O$/DMSO + 0.1% FA), followed by column conditioning and equilibration. Eluted peptides were analysed by the mass spectrometer in positive polarity using a data-independent acquisition mode as follows: an initial MS1 scan was carried out at 120,000 resolution with an AGC target of 3e6 for a maximum IT of 200ms, m/z range: 350-1650. This was followed by sequential MS2 acquisition and fragmentation of ions at 30,000 resolution over 26 variable windows. AGC target set to 3e6 with maximum IT on auto. Normalised collision energy was set to 27%. Total run acquisition time was 76 minutes.

**Mass spectrometry raw data processing and visualisation**

ChIP-MS. Data were processed using the MaxQuant software platform (v1.6.10.43, Tyanova et al)[29], with database searches carried out by the in-built Andromeda search engine against the Swissprot mus musculus database (downloaded – 20230504, entries – 21,965). A reverse decoy database approach was used at a 1% false discovery rate (FDR) for peptide spectrum matches. Search parameters were as follows: maximum missed cleavages set to 2, fixed modification of cysteine carbamidomethylation and variable modifications of methionine oxidation, protein N-terminal acetylation, asparagine deamidation and cyclisation of glutamine to pyro-glutamate. Label-free quantification was enabled with an LFQ minimum ratio count of 1. ‘Match between runs’ function was used with match and alignment time limits of 0.7 and 20 minutes respectively. Reporting of iBAQ values was enabled.

AP-MS. Data were processed using the Spectronaut software platform (Biognosys, v18.5.231110.55695) [30]. Analysis was carried out in direct DIA mode as follows: 1. ***Pulsar Search***: library generation and database search carried out using default settings for a tryspin/p specific digest as follows - missed cleavage rate set to 3 and variable modifications allowed for methionine oxidation, protein N-terminal acetylation, asparagine deamidation and cyclisation of glutamine to pyro-glutamate. PSM, Peptide and Protein group FDR = 0.01. Searches were carried out against the Swissprot mus musculus protein sequence database concatenated with truncated sequence of IKZF1_MOUSE (Uniprot ID: Q03267, downloaded 31/012023, 21,967 entries). Truncated sequence determined by Clustal Omega multi-sequence alignment of canonical sequence and IKZF1 variants. Longest stretch of common sequence, concatenated into database as IKZF1 entry. We reasoned that this would help determine more accurately the relative level of IZF1 across conditions, as quantification would be determined from shared tryptic peptides only. Additionally, searches were also carried against a universal protein contaminants database[28] (downloaded 20220604, 381 entries). 2. ***Direct DIA analysis***: a mutated decoy database approach was employed with protein q-value cut-off for the experiment set to 0.01 at the identification level. Quantification set to MS2 with proteotypicity filter set to only protein group specific with no value imputation strategy employed. Protein quantification method set to MaxLFQ[31]. Two different sets of Spectronaut analyses were carried out. First all samples were analysed together for specific comparisons against the control condition. In this case, raw protein intensities were exported from Spectronaut and used in downstream analysis. Secondly, conditions WT, ED and P were analysed together and normalised (default local normalization option) protein intensity tables were exported for downstream analysis.

Exploratory and technical analysis of data carried out using in-house developed python pipeline with various visualisations using both the Plotly (Available from: https://plotly.com/) and Pandas (https://doi.org/10.5281/zenodo.3509134) plotting libraries. Spectronaut protein level output tables were filtered to remove hits to the contaminants database.
Further analysis and statistical testing carried out using the Perseus platform[32] (v1.6.15.0). Mouse ontology annotations GOBP, GOMF, GOCC and KEGG (downloaded through Perseus GO annotation tool) were assigned by Uniprot protein accessions. Data was log2 transformed before additional filtering and statistical testing. For multi-group comparisons, data was filtered for proteins with >=3 replicate intensities per experimental group. Protein intensities of 1-way ANOVA significant hits (multiple testing correction = permutation-based FDR, threshold = 0.05) were then z-scored in order to HCA-heatmap cluster the data. HCA row cluster ID's were assigned manually and subsequently used to carry out GO enrichment analysis (Fishers Exact test, multiple-testing correction = Benjamini-Hochberg, FDR threshold = 0.05). For 2-group comparisons, intensities were filtered as previously described and Student's t-test carried out (multiple testing correction = permutation-based FDR, threshold = 0.05). T-test results visualised as volcano plots. Intensities of significant hits were z-scored and further processed for GO enrichment analysis as previously described.

### Ikaros motif count and density

Ikaros motif count was calculated as the number of the core Ikaros motifs GGAA, TTCC, GGGA, and TCCC per ATAC-seq peak. Ikaros motif density for each region was calculated as counts/length of the ATAC-seq peak in basepairs.

### GSEA

Gene set enrichment analysis was performed for early Ikaros induced transcription changes (2h vs. 0h) against the following gene sets: reactome integrin cell surface interaction, reactome cell cycle, downregulated in DN Ik6 B-ALL vs B-ALL, and upregulated in DN Ik6 B-ALL vs B-ALL (DE genes of padj<0.01 and FC>1.5 obtained from Supplementary Table NIHMS712753-supplement-4 Tab 3 in Churchman et al 2015[33] )

### References


1 Nesterova, T. B. *et al.* Systematic allelic analysis defines the interplay of key pathways in X chromosome inactivation. *Nat Commun* **10**, 3129, doi:10.1038/s41467-019-11171-3 (2019).

2 Gu, Z., Eils, R. & Schlesner, M. Complex heatmaps reveal patterns and correlations in multidimensional genomic data. *Bioinformatics* **32**, 2847-2849, doi:10.1093/bioinformatics/btw313 (2016).

3 Young, M. D., Wakefield, M. J., Smyth, G. K. & Oshlack, A. Gene ontology analysis for RNA-seq: accounting for selection bias. *Genome Biol* **11**, R14, doi:10.1186/gb-2010-11-2-r14 (2010).

4 Ewels, P. A. *et al.* The nf-core framework for community-curated bioinformatics pipelines. *Nat Biotechnol* **38**, 276-278, doi:10.1038/s41587-020-0439-x (2020).

5 Li, H. & Durbin, R. Fast and accurate short read alignment with Burrows-Wheeler transform. *Bioinformatics* **25**, 1754-1760, doi:10.1093/bioinformatics/btp324 (2009).

6 Ramirez, F. *et al.* deepTools2: a next generation web server for deep-sequencing data analysis. *Nucleic Acids Res* **44**, W160-165, doi:10.1093/nar/gkw257 (2016).

7 Zhang, Y. *et al.* Model-based analysis of ChIP-Seq (MACS). *Genome Biol* **9**, R137, doi:10.1186/gb-2008-9-9-r137 (2008).

8 Amemiya, H. M., Kundaje, A. & Boyle, A. P. The ENCODE Blacklist: Identification of Problematic Regions of the Genome. *Sci Rep* **9**, 9354, doi:10.1038/s41598-019-45839-z (2019).

9 Consortium, E. P. An integrated encyclopedia of DNA elements in the human genome. *Nature* **489**, 57-74, doi:10.1038/nature11247 (2012).

10 Liao, Y., Smyth, G. K. & Shi, W. The R package Rsubread is easier, faster, cheaper and better for alignment and quantification of RNA sequencing reads. *Nucleic Acids Res* **47**, e47, doi:10.1093/nar/gkz114 (2019).

11 Love, M. I., Huber, W. & Anders, S. Moderated estimation of fold change and dispersion for RNA-seq data with DESeq2. *Genome Biol* **15**, 550, doi:10.1186/s13059-014-0550-8 (2014).

12 Yu, G., Wang, L. G. & He, Q. Y. ChIPseeker: an R/Bioconductor package for ChIP peak annotation, comparison and visualization. *Bioinformatics* **31**, 2382-2383, doi:10.1093/bioinformatics/btv145 (2015).

13 Danecek, P. *et al.* Twelve years of SAMtools and BCFtools. *Gigascience* **10**, doi:10.1093/gigascience/giab008 (2021).

14 Quinlan, A. R. & Hall, I. M. BEDTools: a flexible suite of utilities for comparing genomic features. *Bioinformatics* **26**, 841-842, doi:10.1093/bioinformatics/btq033 (2010).

15 Zhang, T. *et al.* A variant NuRD complex containing PWWP2A/B excludes MBD2/3 to regulate transcription at active genes. *Nat Commun* **9**, 3798, doi:10.1038/s41467-018-06235-9 (2018).

16 Fursova, N. A. *et al.* Synergy between Variant PRC1 Complexes Defines Polycomb-Mediated Gene Repression. *Mol Cell* **74**, 1020-1036 e1028, doi:10.1016/j.molcel.2019.03.024 (2019).

17 Langmead, B. & Salzberg, S. L. Fast gapped-read alignment with Bowtie 2. *Nat Methods* **9**, 357-359, doi:10.1038/nmeth.1923 (2012).

18 Lawrence, M. *et al.* Software for computing and annotating genomic ranges. *PLoS Comput Biol* **9**, e1003118, doi:10.1371/journal.pcbi.1003118 (2013).
19 Koohy, H. *et al.* Genome organization and chromatin analysis identify transcriptional downregulation of insulin-like growth factor signaling as a hallmark of aging in developing B cells. *Genome Biol* **19**, 126, doi:10.1186/s13059-018-1489-y (2018).
20 Rao, S. S. *et al.* A 3D map of the human genome at kilobase resolution reveals principles of chromatin looping. *Cell* **159**, 1665-1680, doi:10.1016/j.cell.2014.11.021 (2014).
21 Loven, J. *et al.* Selective inhibition of tumor oncogenes by disruption of super-enhancers. *Cell* **153**, 320-334, doi:10.1016/j.cell.2013.03.036 (2013).
22 Whyte, W. A. *et al.* Master transcription factors and mediator establish super-enhancers at key cell identity genes. *Cell* **153**, 307-319, doi:10.1016/j.cell.2013.03.035 (2013).
23 Consortium, F. *et al.* A promoter-level mammalian expression atlas. *Nature* **507**, 462-470, doi:10.1038/nature13182 (2014).
24 Bonev, B. *et al.* Multiscale 3D Genome Rewiring during Mouse Neural Development. *Cell* **171**, 557-572 e524, doi:10.1016/j.cell.2017.09.043 (2017).
25 Hill, L. *et al.* Igh and Igk loci use different folding principles for V gene recombination due to distinct chromosomal architectures of pro-B and pre-B cells. *Nat Commun* **14**, 2316, doi:10.1038/s41467-023-37994-9 (2023).
26 Hsieh, T. S. *et al.* Resolving the 3D Landscape of Transcription-Linked Mammalian Chromatin Folding. *Mol Cell* **78**, 539-553 e538, doi:10.1016/j.molcel.2020.03.002 (2020).
27 Mohammed, H. *et al.* Rapid immunoprecipitation mass spectrometry of endogenous proteins (RIME) for analysis of chromatin complexes. *Nat Protoc* **11**, 316-326, doi:10.1038/nprot.2016.020 (2016).
28 Turriziani, B. *et al.* On-beads digestion in conjunction with data-dependent mass spectrometry: a shortcut to quantitative and dynamic interaction proteomics. *Biology (Basel)* **3**, 320-332, doi:10.3390/biology3020320 (2014).
29 Tyanova, S., Temu, T. & Cox, J. The MaxQuant computational platform for mass spectrometry-based shotgun proteomics. *Nat Protoc* **11**, 2301-2319, doi:10.1038/nprot.2016.136 (2016).
30 Bruderer, R. *et al.* Extending the limits of quantitative proteome profiling with data-independent acquisition and application to acetaminophen-treated three-dimensional liver microtissues. *Mol Cell Proteomics* **14**, 1400-1410, doi:10.1074/mcp.M114.044305 (2015).
31 Cox, J. *et al.* Accurate proteome-wide label-free quantification by delayed normalization and maximal peptide ratio extraction, termed MaxLFQ. *Mol Cell Proteomics* **13**, 2513-2526, doi:10.1074/mcp.M113.031591 (2014).
32 Tyanova, S. *et al.* The Perseus computational platform for comprehensive analysis of (prote)omics data. *Nat Methods* **13**, 731-740, doi:10.1038/nmeth.3901 (2016).
33 Churchman, M. L. *et al.* Efficacy of Retinoids in IKZF1-Mutated BCR-ABL1 Acute Lymphoblastic Leukemia. *Cancer Cell* **28**, 343-356, doi:10.1016/j.ccell.2015.07.016 (2015).
